# Targeted disruption of linkage-specific ubiquitylation reveals a key role of K29-linked ubiquitylation in epigenome integrity

**DOI:** 10.1101/2024.10.29.620783

**Authors:** Javier Arroyo-Gomez, Juanjuan Wang, Claire Guérillon, Ekaterina Isaakova, Nazaret Reverón-Gómez, Mikaela Koutrouli, Aldwin Suryo Rahmanto, Katrine Mitrofanov, Andreas Ingham, Sofie Schovsbo, Katrine Weischenfeldt, Fabian Coscia, Dimitris Typas, Moritz Völker-Albert, Victor Solis, Lars Juhl Jensen, Anja Groth, Andreas Mund, Petra Beli, Robert F Shearer, Niels Mailand

## Abstract

Linkage-specific ubiquitin chains dictate the functional outcome of numerous critical ubiquitin-dependent signaling processes. However, the functions and targets of several poly-ubiquitin topologies remain poorly defined due to a paucity of tools for their specific detection and manipulation. To remedy this knowledge gap, we applied a cell-based ubiquitin replacement strategy enabling targeted conditional abrogation of each of the seven lysine-based ubiquitin chain types in human cells to profile system-wide impacts of disabling formation of individual chain types. Focusing on K29-linked ubiquitylation, we reveal a strong association of this linkage type with chromatin-associated proteins and show that the H3K9me3 methyltransferase SUV39H1 is a prominent cellular target of this modification. We demonstrate that K29-linked ubiquitylation is essential for proteasomal degradation of SUV39H1 despite its extensive modification by K48-linked ubiquitylation, and that K29-linked ubiquitylation of SUV39H1 is catalyzed and reversed by TRIP12 and TRABID, respectively. Preventing K29-linked ubiquitylation-mediated control of SUV39H1 stability deregulates the H3K9me3 landscape, but not other histone marks. Collectively, our ubiquitin replacement cell line panel and datasets provide valuable resources for illuminating cellular functions of linkage-specific ubiquitin chains and establish a key role of K29-linked ubiquitylation in epigenome integrity.

## Introduction

Cellular homeostasis is maintained through a highly regulated balance between protein production and degradation, the latter of which is extensively mediated through the ubiquitin-proteasome system. Ubiquitin (Ub) is a highly abundant small modifier protein that is attached to substrate proteins through a hierarchical ubiquitylation cascade. Ub is activated in an ATP-dependent manner by an E1 Ub-activating enzyme and transferred to an E2 Ub-conjugating enzyme before it is covalently attached to an acceptor lysine on a substrate protein by an E3 Ub ligase ^1^. Protein ubiquitylation is a reversible process that is antagonized by the action of deubiquitinases (DUBs), providing versatility to the Ub signaling process ^2^. Over 600 known E3 Ub ligase enzymes ^3^ and approximately 100 DUBs ^2^ are encoded by human cells, conferring a high degree of specificity to the Ub system. Attachment of a single Ub moiety (mono-Ub) can elicit structural changes or modulate interaction surfaces of a substrate protein. However, the versatility of Ub signaling outcomes arises from chain extension via one of seven internal lysine residues (K6, K11, K27, K29, K33, K48 and K63), or the N-terminal Methionine (M1), to form structurally distinct Ub polymers (poly-Ub). Chain assembly can also occur through multiple Ub-linkage sites simultaneously, giving rise to heterogeneous or branched chains, expanding Ub-chain complexity ^1^. Furthermore, recent findings have indicated that Ub can be attached to non-lysine amino acids and even to unconventional acceptor molecules such as lipids and sugars ^4^.

The most common outcome of Ub signaling is proteasomal degradation via the 26S proteasome, to which substrates are directed when modified with K48-linked Ub chains ^5^. However, other Ub-linkages are associated with degradation in certain contexts. For example, K29-linked Ub chains are heavily upregulated during proteotoxic stress, colocalize with stress granule components, and enhance degradation signaling through facilitating p97/VCP-mediated unfolding ^6^, which is primarily required to extract degradation substrates embedded in macromolecular structures such as chromatin ^7^. More recently, K6-linked Ub chains driven by RNF14 were implicated in proteasome- and p97-dependent resolution of RNA-protein crosslinks ^8,9^. Ub chains can mediate receptor interactions rather than signal degradation by forming a binding surface for so-called Ub “readers”. Although many Ub interactions are likely linkage-independent, it is expected that most Ub-linkage types have specific interaction partners as a proteome-wide study probing with synthetic di-Ub found many linkage-specific interactions ^10^. Ub-mediated interactions are conferred through mostly alpha-helical structures forming a diverse range of Ub binding domains with various degrees of interaction affinity or Ub-linkage specificity ^11^, a classic example of which can be seen in DNA repair. Upon the formation of DNA double-strand breaks (DSBs), K63- and K27-linked Ub chains, catalyzed by RNF8-UBC13 and RNF168 respectively, form at DSB sites providing a scaffold for the recruitment of downstream DSB repair factors ^12,13^.

The heterogeneous and dynamic nature of Ub signaling forms an important cellular code; however, not all Ub-linkage types are well characterized. Under normal cycling conditions, M1-, K6-, K27-, and K33-linked Ub chains are found in low abundance in mammalian cells (usually undetectable or <0.5%) ^14^. The function of these atypical Ub-linkage types has been explored; however, due to technical limitations, broad biological functions have yet to be resolved. Furthermore, current understanding of low abundance Ub-linkages often relies upon artefact-prone Ub mutant overexpression that may alter endogenous Ub signaling. Proteomic approaches allow quantitative measurement of Ub-linkage composition on individual substrates; however, methodologies for the high throughput study of Ub-linkages in a complex mixture are lacking. This is largely due to the difficulty in developing robust enrichment reagents for multiple Ub-linkage types.

To address this limitation, we leveraged the Ub-replacement strategy, a conditional cell-based system for depletion of the endogenous Ub pool that is rescued to near-endogenous levels by coexpression of exogenous Ub harboring selected K-to-R mutations ^15^. We previously utilized this Ub-replacement approach to show that K27-linked Ub chains are essential for fitness of mammalian cells, predominantly nuclear and are associated with p97 activity in the nucleus ^16^. Here, we expand upon the cellular Ub replacement resource to profile the landscape of lysine-based Ub substrates in a linkage-specific manner for a mammalian cancer cell line model. Among other proteins, we identified the histone methyltransferase SUV39H1 as a prominent substrate of proteolytic K29-linked ubiquitylation. Stabilization of SUV39H1 by abrogation of K29-Ub chain formation increases the SUV39H1-mediated epigenetic modification H3K9me3 ^17^ that governs heterochromatin formation ^18^. Furthermore, we identify the HECT E3 Ub ligase TRIP12 and the DUB TRABID (aka ZRANB1) as the major writer and eraser of nuclear K29-linked ubiquitylation and thus SUV39H1 levels. Furthermore, deregulation of this E3-DUB signalling pathway is prevalent in lung cancer, and is associated with a poorer prognosis in multiple cancer types. Taken together, our findings highlight a key role for K29-linked ubiquitylation in epigenome integrity that may have implications for human disease.

## Results

### An expanded Ub replacement system for conditional abrogation of individual lysine-based Ub chains

We expanded upon the previously described Ub replacement strategy ^15^ to include all seven structurally diverse lysine-based linkages (**Figure 1A**). *In silico* folding prediction of full-length Ub was used to determine structural strain (S_i_) resulting from individual lysine to arginine mutations (K-to-R), which was negligible (**Figure 1A**). It should be noted that the S_i_ for the Ub K27R mutation, although low, was relatively higher than other Ub K-to-R mutations (**Figure 1A**), however we previously showed that the Ub K27R mutation is fully permissive for Ub conjugation in cells ^16^. We next conducted a series of two-step stable transfections of a U2OS human osteosarcoma cell line to generate a panel of doxycycline-inducible Ub replacement cell lines for conditional abrogation of each individual linkage type (**Figure 1B,C**). We had previously generated a stable U2OS cell line harboring a cassette encoding inducible shRNAs targeting the four human loci containing Ub-coding genes (herein U2OS/shUb) ^16^. We utilized the U2OS/shUb base to generate a series of derivative cell lines harboring a cassette encoding human fusion proteins UBA52 and RPS27A with Ub either in wild-type (WT) or K-to-R form (herein U2OS/shUb/HA-Ub(WT) or U2OS/shUb/HA-Ub(K-to-R)) (**Figure 1B,C**). Successful induction of Ub replacement was determined by detection of conjugated Ub after doxycycline treatment via immunofluorescence and immunoblot analysis. Stable clones were carefully selected for optimal Ub expression levels, both in uniformity and similarity to endogenous Ub levels, and doxycycline-induced Ub replacement in these cell lines was validated by qPCR analysis (**Figure 1D-G**; **Figure S1A,B**). A characteristic Ub smear was visible by immunoblot for all Ub replaced cell lines indicating functional Ub-polymer formation (**Figure S1A**), and proteasomal degradation was inhibited only in the U2OS/shUb/HA-Ub(K48R) replaced cell line when induced with a characterized proteolysis targeting chimera (PROTAC) driving VHL-dependent degradation of BRD4 ^19^, indicating proteolytic function of the Ub K-to-R mutants was unimpaired except for the expected impact of abrogating K48-linked Ub chain formation (**Figure S1C**). To validate normal function of replaced Ub in the context of DNA repair, we induced the DSB response in the Ub replacement cell line panel by treatment with ionizing radiation (IR), and measured 53BP1 foci formation, a Ub-dependent readout of canonical DSB repair ^20^. Recruitment of 53BP1 to damage sites was unimpaired except in Ub(K63R)- and Ub(K27R)-replaced cells or when Ub signaling was inhibited with an E1 Ub inhibitor (**Figure 1H,I**), consistent with the known requirement for both K63- and K27-linked Ub for efficient 53BP1 recruitment to DSBs ^12,13^. Moreover, we used Ub(K6R) replacement cells and a K6-Ub-specific binder (LotA) to verify recent findings showing a strong increase in K6-linked ubiquitylation upon RNA-protein crosslink induction by combined UV-A and 4-thiouridine (S4U) treatment ^8^ (**Figure 1J**; **Figure S1D**).

**Figure 1.**
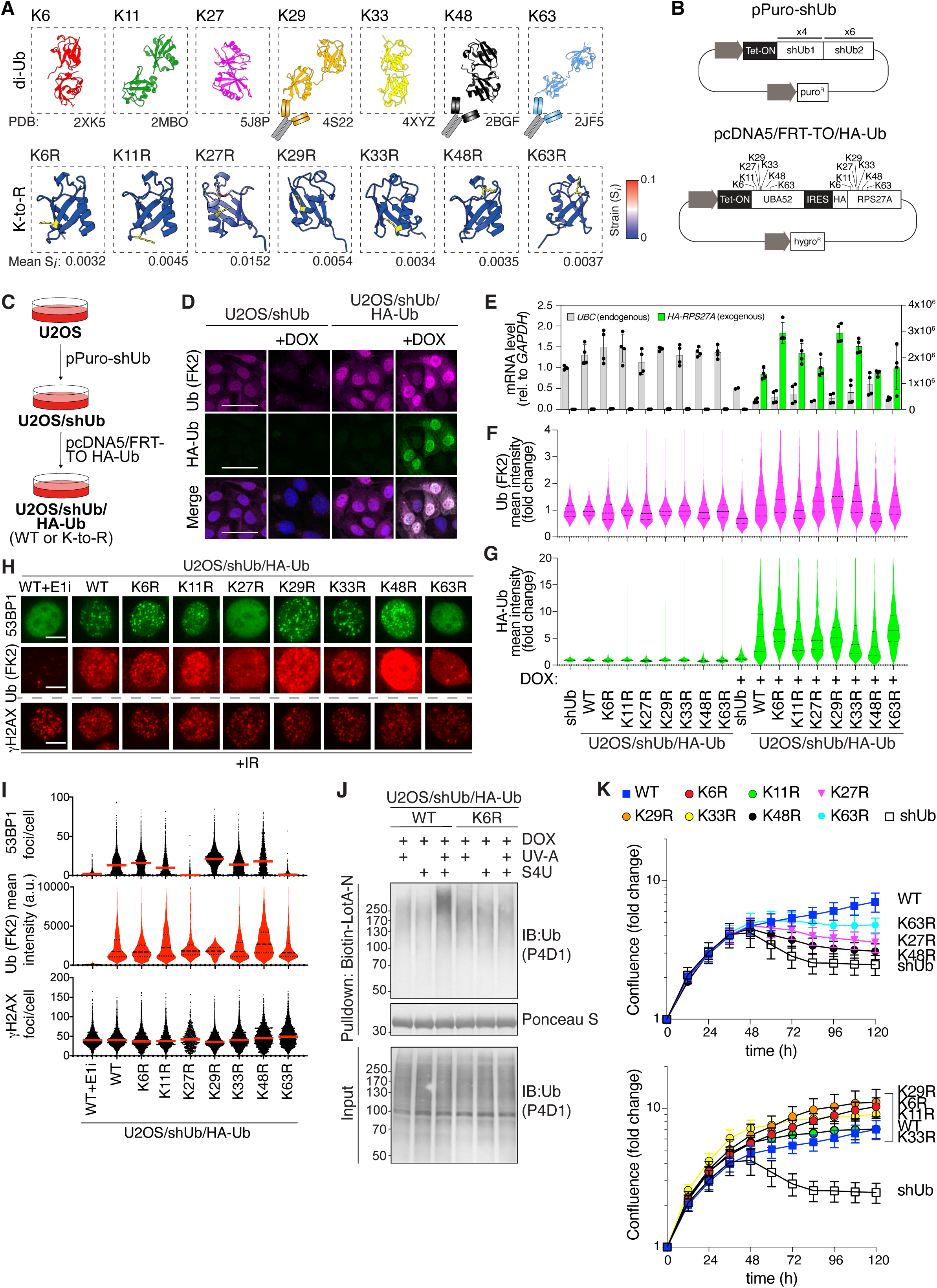
A Ub replacement cell line panel for inducible abrogation of lysine-based Ub topologies. **A.** Schematic depicting structures of lysine linked di-Ub (upper panel). PDB identifiers are indicated. Summary of testing of commercially available Ub enrichment reagents with linkage specificity by immunoblot (IB) and immunofluorescence (IF) is indicated (see **Figure S1C**). Alphafold2-based prediction (lower panel) of structural strain of Ub as a result of indicated K-to-R mutations (highlighted in yellow), expressed as Strain per residue (S_i_). Strain on WT structure calculated from average of 5 output structures. **B.** Schematic of Doxycycline (DOX)-inducible pPuro-shUb and pcDNA5/FRT-TO/HA-Ub expression plasmids required for two-step Ub replacement (C). **C.** Stepwise selection procedure for generation of DOX-inducible Ub replacement U2OS cell lines. **D.** Representative images of U2OS/shUb and U2OS/shUb/HA-Ub(WT) derivative U2OS cell lines that were treated or not with DOX for 48 or 72 h, respectively, and immunostained with indicated antibodies. FK2 staining indicates conjugated Ub. Scale bar, 50 µm. **E.** U2OS/shUb and derivative replacement cell lines were treated or not with DOX and mRNA levels were analyzed by RT-qPCR. Primers to GAPDH were used as a normalization control (data are technical duplicates of two independent experiments). Left axis corresponds to *UBC* detection and right axis corresponds to *HA-RPS27A* detection. **F,G**. Quantitation of conjugated Ub (FK2) (F) and HA (G) immunostaining of U2OS/shUb and derivative replacement cell lines treated or not with DOX after DOX treatment for the indicated times, determined by quantitative image-based cytometry (QIBC) (thick dashed-line, median; dotted-lines, quartiles). Data is a single representative replicate from 3 independent experiments, expressed as fold change normalized to untreated cells (>1000 cells analyzed per sample). **H**. Representative images of DOX-treated Ub replacement cell lines exposed to ionizing radiation (IR, 2 Gy) 1 h prior to CSK pre-extraction, fixation and immunostaining with indicated antibodies. Scale bar, 10 µm. **I**. Quantitation of 53BP1 (upper), FK2 (middle) and ψH2AX (lower panel) staining localized to IR-induced foci in cells in (H), using QIBC (thick dashed-line, median; dotted-lines, quartiles). Data is a single representative replicate from 3 independent experiments (>1500 cells analyzed per condition). **J**. Immunoblot (IB) analysis of K6-linked Ub chains isolated by pulldown with the K6-Ub binder Biotin-LotA-N in the indicated DOX-treated Ub replacement cell lines exposed to UV-A (500 mJ/cm^2^) and/or 4-thiouridine (S4U, 25 mM), combined treatment of which leads to RNA-protein crosslink formation. **K**. Short-term proliferation of U2OS/shUb and derivative Ub replacement cell lines after DOX induction for the indicated times. Cell lines that proliferate in a manner comparable to WT include K29R, K6R, K11R and K33R replaced U2OS cells (lower), while K63R, K27R, K48R and shUb-only replaced cell lines proliferated sub-optimally (upper panel) after DOX addition. Normalized logarithmic proliferation was determined using Incucyte image-based confluence assay (mean±s.e.m.; *n*=5 independent experiments).

The relatively poor characterization of roles for low abundance Ub chains is in part due to the limited range of robust linkage-specific Ub detection reagents. For this reason, the Ub replacement cell line panel was used to assess the performance and specificity of commercially available Ub linkage-selective enrichment reagents for immunoblotting and immunofluorescence. In our hands, a range of such affinity reagents displayed sub-optimal linkage specificity when utilized by a standard immunofluorescence protocol, except for the recently described sAB-K29 binder ^6^ and the APU2 antibody recognizing K48-linked chains (**Figure S1E**). An antibody recognizing K63-linked chains performed well by immunoblot but showed non-specific staining by imaging (**Figure S1E**). The panel of conditional Ub K-to-R replacement cell lines thus provides a useful tool for assessing the specificity of linkage-selective Ub affinity reagents and the requirement of individual Ub chain types for cellular processes of interest.

### High abundance Ub chains are required for efficient cell proliferation

We next utilized the complete Ub replacement panel to assess the importance of individual Ub topologies for mammalian cell viability. As expected, abrogation of high abundance Ub topologies (K48- and K63-linked Ub) resulted in poor short-term viability, as assessed by relative cellular confluence (**Figure 1K**). Reduced cellular viability observed in Ub(K48R)- and Ub(K63R)-replaced cells was accompanied by an increased proportion of cells in G2/M phase and a reduced fraction of mitotic cells, indicative of G2 arrest (**Figure S1F**). Loss of several low abundance Ub topologies (K6-, K11- and K33-linked Ub) had minor impacts on cell proliferation in short term confluence assays (**Figure 1K**). As previously reported ^16^, cells with loss of K27-linked Ub showed poor viability, while abrogation of K29-linked ubiquitylation was well tolerated.

### A global map of ubiquitylation changes resulting from loss of individual Ub chain types

Given the lack of functional understanding of low abundance lysine-based Ub topologies in cell biology, we used our panel of Ub replacement cell lines to profile global changes in the Ub substrate landscape of cells with loss of individual Ub chain types. To this end, we subjected HA-Ub conjugates isolated from Ub replacement cell lines under denaturing conditions and corresponding whole cell extracts following 72 h of Ub replacement to quantitative label-free MS proteomic analysis, and individual lysine mutant samples were compared to samples from Ub(WT)-replaced cells (**Figure 2A**). Among Ub-replaced cell lines, the most pronounced impacts on whole proteome status were observed for abrogation of K48- and K63-linked ubiquitylation, consistent with these chains comprising the most abundant Ub-linkage types in cells, while disrupting other Ub chain types had more modest effects (**Figure S2A; Dataset S1**). Due to the particularly strong adverse impact of Ub(K48R) replacement on cell proliferation (**Figure 1K**), we examined whole proteome impacts of Ub(K48R)-replaced cell lines at both 48 and 72 h time points, the latter of which displayed more extensive proteomic changes (**Figure S2A**).

**Figure 2.**
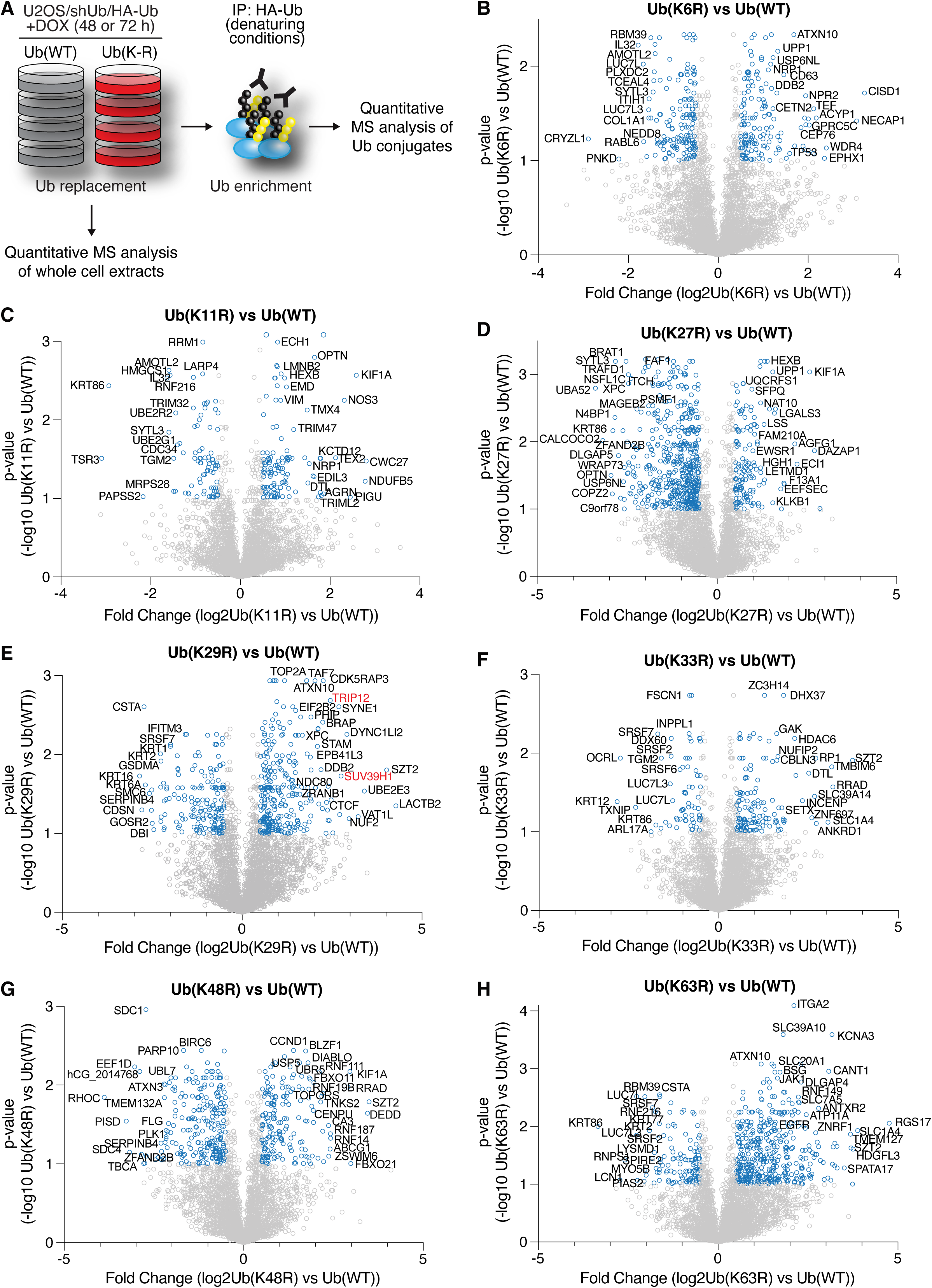
Global ubiquitylation changes induced by disrupting formation of individual Ub chain types. **A.** Schematic depicting experimental setup for proteomic analysis of HA-Ub conjugates enriched from whole cell lysates from Ub(WT) and Ub(K-to-R) replacement cell lines after DOX treatment for 48 or 72 h. **B**-**H**. Ubiquitylation changes resulting from replacement of endogenous Ub with indicated Ub(K-to-R) mutants by DOX treatment for 72 h, analyzed by quantitative label-free MS as shown in (A). Volcano plots depict protein abundance changes after denaturing HA-enrichment in HA-Ub(K-to-R)-replaced cells relative to HA-Ub(WT)-replaced cells. Blue dots indicate proteins with at least 1.5-fold enrichment and passing a significance threshold of p<0.1. The most strongly regulated proteins are labelled. See **Table S2** for full results.

We then performed pairwise comparisons between HA-Ub conjugates isolated from Ub(WT)- and Ub(K-to-R)-replaced cells to profile global ubiquitylation changes induced by abrogation of individual Ub chain linkages (**Figure 2B-H**; **Dataset S2**). Principal component analysis (PCA) showed expected clustering of proteomic samples (**Figure S2B**). The ubiquitylation profile of K27R-replaced cells matched that of our previously described results ^16^, including decreased overall ubiquitylation of multiple p97/VCP cofactors including UFD1, NSFL1C/p47 and FAF1, as well as the nucleotide excision repair factors RAD23B and XPC. We used gene ontology (GO) term analysis applied to the ubiquitylation datasets to pinpoint cellular processes responsive to disruption of specific Ub linkage topologies. For instance, Ub(K63R)-replaced cells uniquely showed significant enrichment of terms related to cell membranes (**Figure S3**), consistent with the well-established role of K63-linked ubiquitylation in membrane trafficking ^21^. Likewise, abrogation of K33-linked ubiquitylation impacted the ubiquitylation status of proteins involved in multiple aspects of RNA processing, whereas disrupting K27-linked ubiquitylation, but not other Ub-linkages, was associated with enhanced ubiquitylation of mitochondrial proteins (**Figure S3**). Thus, ubiquitylation changes arising from abolishing formation of specific Ub chain types shed light on the nature of cellular processes regulated by these Ub-linkage topologies. Notably, Ub(K29R)-replaced samples contrasted to other Ub(K-to-R) mutants in that enriching proteins were often nuclear localizing; consistently GO term analysis revealed a striking enrichment of chromosome-associated processes that was selective to this Ub linkage type (**Figure 3A**; **Figure S3**), which we set out to explore further. Among others, notable proteins enriched in Ub(K29R)-replaced samples included E3 ubiquitin-protein ligases TRIP12 and RNF168 (**Figure 2E**). In line with this, TRIP12 is known to catalyze predominantly K29-linked Ub chains *in vitro*, and often in branching complexes with K48-linked Ub ^22^.

**Figure 3.**
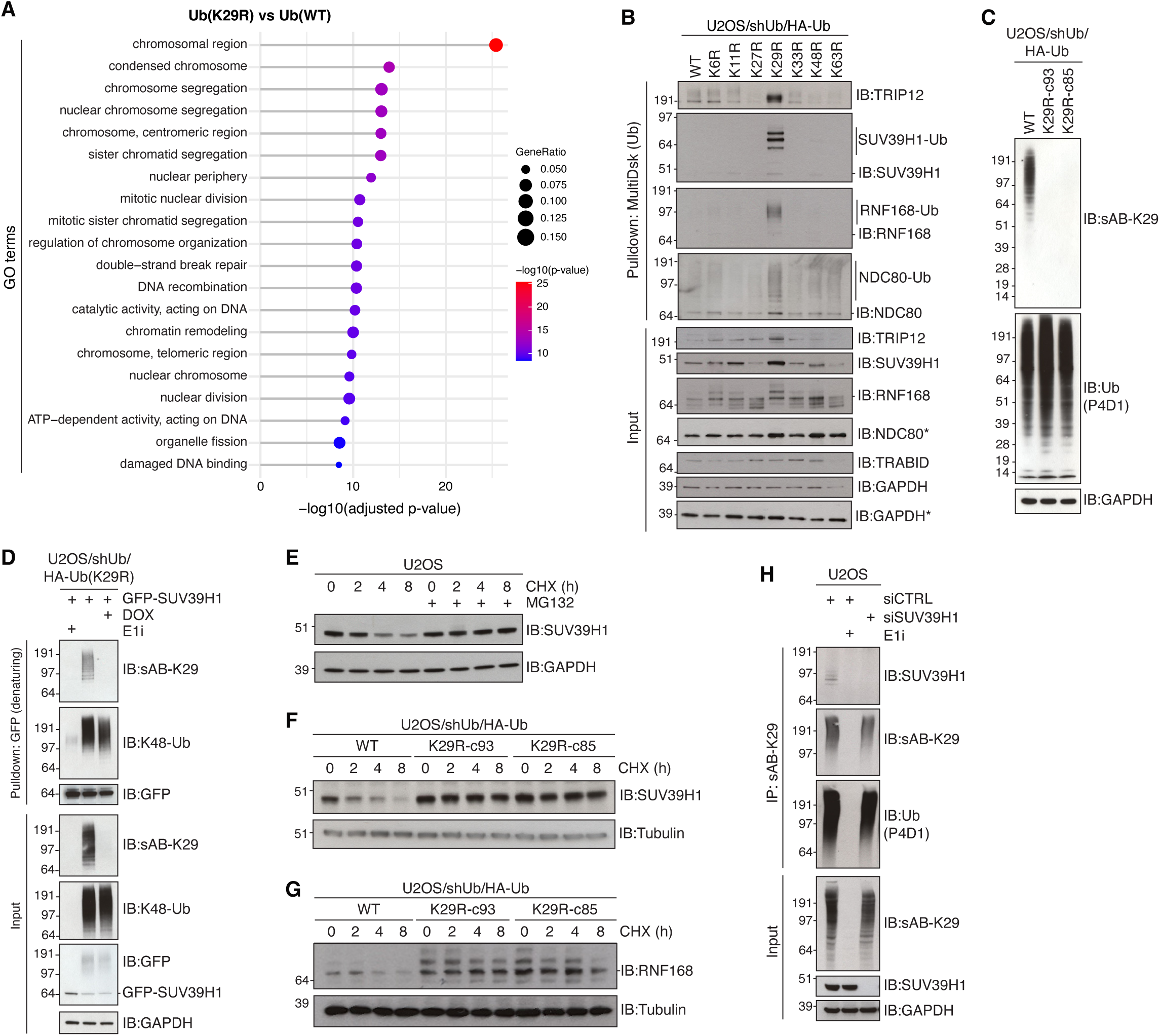
K29-linked ubiquitylation of the H3K9me3 methyltransferase SUV39H1 is required for its proteasomal degradation. **A.** GO term analysis of cellular compartments enriched among proteins showing significantly elevated ubiquitylation in Ub(K29R)-replaced cells relative to Ub(WT)-replaced cells (**Figure 2E**). **B.** Immunoblot analysis of Ub conjugates from indicated Ub-replaced cell lines isolated via MultiDsk pulldown. **C.** Immunoblot analysis of indicated DOX-treated Ub replacement cell lines using sAB-K29 and total Ub (P4D1) antibody. **D.** Ub(K29R) cells transfected with GFP-SUV39H1 expression constructs and treated with DOX or Ub E1 inhibitor (E1i) as indicated were subjected to GFP pulldown under denaturing conditions and immunoblotted with indicated antibodies. **E.** Immunoblot analysis of SUV39H1 in U2OS cells treated or not with cycloheximide (CHX) for the indicated times in the absence or presence of the proteasome inhibitor MG132. **F.** As in (E), using indicated DOX-treated Ub replacement cell lines. **G.** As in (F), but using RNF168 antibody. **H.** U2OS cells transfected with non-targeting control (CTRL) or SUV39H1 siRNAs and treated or not with Ub E1 inhibitor were subjected to immunoprecipitation (IP) with sAB-K29 and immunoblotted with indicated antibodies.

Furthermore, RNF168 is a known proteolytic substrate of TRIP12 under normal cycling conditions ^23^. Among other strongly enriched chromatin-associated proteins was SUV39H1, a histone methyltransferase that catalyzes the formation of histone H3 lysine 9 trimethylation (H3K9me3) in chromatin ^17^ and kinetochore complex component NDC80, a central component of the NDC80 complex which forms at the contact point between kinetochores and microtubules ^24^ (**Figure 2E**). Strikingly, when Ub conjugates enriched from Ub(K29R)-replaced cells were immunoblotted using antibodies to TRIP12, SUV39H1, RNF168 and NDC80, all proteins displayed a high intensity molecular weight shift consistent with covalent poly-Ub modification exclusively in Ub(K29R)-replaced cells (**Figure 3B**). This suggests that the turnover of Ub-modified forms of these proteins becomes defective when formation of K29-linkages is blocked. Indeed, all of these proteins displayed markedly elevated total abundance upon abrogation of K29-linked Ub chains, but not other linkage types (**Figure 3B**). These analyses provide a comprehensive picture of the impact of abrogating specific Ub-linkage types on global ubiquitylation dynamics in human cells, informing on cellular processes regulated by these Ub modifications.

### SUV39H1 is a substrate of proteasomal degradation via K29-linked Ub

SUV39H1 plays a central role in epigenetic maintenance ^17,18^, however how histone methyl transferases and other epigenetic modifiers are regulated remains poorly understood. For this reason, we sought to understand the emerging role of K29-linked Ub chains in regulating SUV39H1. For validation purposes an independent Ub(K29R) replacement clone was generated that showed similar Ub replacement kinetics (**Figure 3C**; **Figure S4A,B**). A synthetic antibody selectively recognizing K29-Ub (sAB-K29 ^6^; **Figure S1C**) was used to validate near-complete abrogation of K29-linked ubiquitylation in these Ub(K29R)-replaced cell lines (**Figure 3C**; **Figure S4B**), and both clones were confirmed to be undergoing correct transcription and translation using EU and L-AHA incorporation, respectively (**Figure S4C**). To confirm the topology of Ub chains formed on SUV39H1, a GFP-SUV39H1 construct was transfected into Ub(K29R)-replaced cells to enable SUV39H1 isolation by GFP immunoprecipitation under denaturing conditions. This showed that K29-linked Ub chains were readily detectable on GFP-SUV39H1 by immunoblotting (**Figure 3D**). Interestingly, K48-linked Ub, which was also robustly detected on GFP-SUV39H1, was still present when K29-linked ubiquitylation was prevented by Ub(K29R) replacement (**Figure 3D**). Detection of K29-and K48-linked Ub chains on both SUV39H1 and RNF168 using linkage-specific antibodies was further observed in parental U2OS cells stably expressing GFP-tagged expression vectors (**Figure 3D**; **Figure S4D**), and a similar profile was observed using a FLAG-SUV39H1 expression construct to validate the Ub signal is not due to ubiquitylation of the GFP-tag (**Figure S4E**). A cycloheximide pulse-chase experiment to assay protein stability confirmed that endogenous SUV39H1 is a substrate of proteasomal degradation with a relatively short half-life as previously reported (**Figure 3E**) ^25^, revealing that SUV39H1 turnover is fully dependent on K29-linked ubiquitylation as evidenced by its complete stabilization in two independent clones of Ub(K29R)-replaced cells (**Figure 3F**). Similar K29-Ub-dependent degradation kinetics were observed for RNF168 (**Figure 3G**). Finally, the presence of K29-linked Ub chains on endogenous SUV39H1 was confirmed by immunoblotting of K29-linked Ub immunoprecipitates (**Figure 3H**). Collectively, these results highlight an important role for K29-linked ubiquitylation in Ub-dependent regulation of chromatin-associated proteins and establish SUV39H1 as a novel substrate of proteasomal degradation via K29-linked Ub.

### TRIP12 is required for K29-linked ubiquitylation of SUV39H1

Due to the enrichment of chromosome-associated proteins affected by K29-linked ubiquitylation, we next sought to explore Ub writer machineries involved in nuclear K29-linked ubiquitylation in an unbiased manner. To this end, we performed a high-content imaging-based siRNA screen for E3 Ub ligases impacting K29-Ub abundance using sAB-K29 binder detection in the nucleus as a readout. The specificity of sAB-K29 in detecting K29-Ub signals by immunostaining was confirmed in two independent Ub(K29R) clones (**Figure S5A**). U2OS cells were transfected with an siRNA library comprehensively covering known E3 Ub ligase enzymes and immunostained with sAB-K29 (**Figure 4A**; **Dataset S3**). Notably, depletion of TRIP12 resulted in the strongest decrease in nuclear K29-Ub detection (**Figure 4B**), with an approximate 50% decrease in nuclear K29-Ub signal relative to a non-targeting siRNA control (**Dataset S3**). Interestingly, a moderate decrease in detectable K29-Ub signal was also observed when depleting several Cullin proteins (CUL1, CUL4A, CUL4B or CUL5), scaffold proteins for Cullin-RING Ub ligases (**Figure 4B**). Reduced detection of nuclear K29-Ub after TRIP12 depletion was also observed in HeLa cells (**Figure S5B,C**).

**Figure 4.**
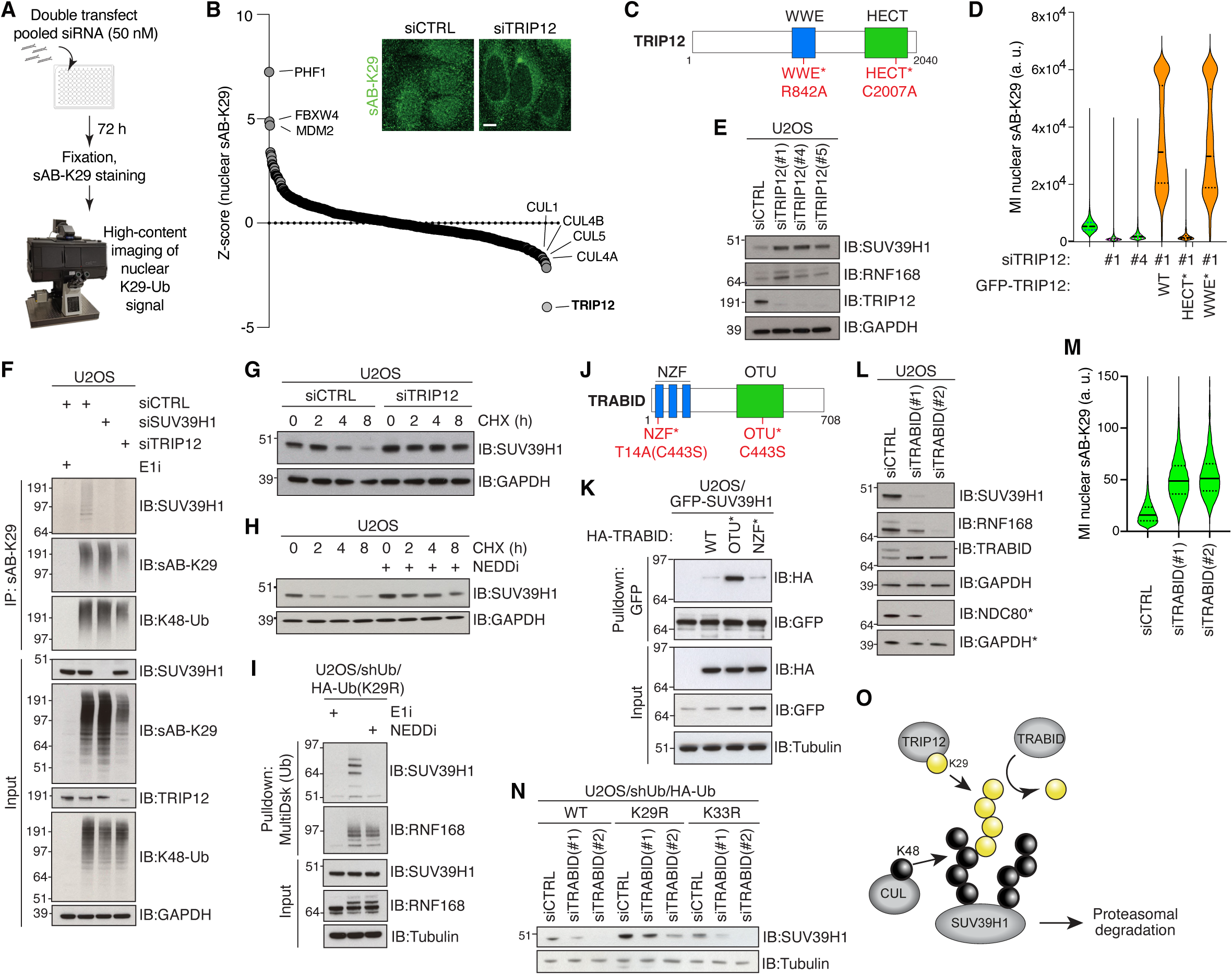
K29-linked ubiquitylation of SUV39H1 is controlled by TRIP12 and TRABID. **A.** Schematic depicting workflow of siRNA screen for regulators of K29-linked ubiquitylation in the nucleus. **B.** Waterfall plot summarizing siRNA screen for regulators of nuclear K29-linked ubiquitylation in U2OS cells (A). Library contains multiplexed siRNAs to 575 E3 Ub ligase enzymes. Data shows Z-score of sAB-K29 immunostaining relative to population by QIBC. See **Table S3** for full results. Inset shows representative images of cells transfected with indicated siRNAs and immunostained with sAB-K29. Scale bar, 10 µm. **C.** Schematic of human TRIP12 protein showing functional domains and mutations introduced to generate indicated mutants. **D.** QIBC analysis of nuclear sAB-K29 immunostaining in U2OS cells sequentially transfected with indicated siRNA and siRNA-resistant GFP-TRIP12 expression plasmids. Samples transfected with GFP-TRIP12 plasmids (orange) were gated on GFP-positive cells. Data is a single representative replicate from 3 independent experiments (thick dashed lines, median; dotted lines, quartiles; >1000 cells analyzed per condition). **E.** Immunoblot analysis of U2OS cells transfected with non-targeting control (CTRL) or TRIP12 siRNAs. **F.** U2OS cells transfected with indicated siRNAs and treated or not with Ub E1 inhibitor were subjected to immunoprecipitation (IP) with sAB-K29 and immunoblotted with indicated antibodies. **G.** Immunoblot analysis of siRNA-transfected U2OS cells treated or not with cycloheximide (CHX) for the indicated times. **H.** Immunoblot analysis of SUV39H1 in U2OS cells treated or not with cycloheximide (CHX) for the indicated times in the absence or presence of a NEDDylation inhibitor (NEDDi). **I.** Immunoblot analysis of Ub conjugates isolated via MultiDsk pulldown from Ub(K29R)-replaced cell lines treated with Ub E1 inhibitor or NEDDylation inhibitor. **J.** Schematic of human TRABID protein showing functional domains and mutations introduced to generate indicated mutants. **K.** U2OS cells stably expressing GFP-SUV39H1 were transfected with indicated HA-TRABID expression constructs and subjected to GFP pulldown, followed by immunoblotting with indicated antibodies. **L.** Immunoblot analysis of U2OS cells transfected with indicated siRNAs. Note that siTRABID(#1) has reduced knockdown efficiency relative to siTRABID(#2) (* indicates samples run on a separate gel). **M.** QIBC analysis of nuclear sAB-K29 immunostaining in U2OS cells treated with indicated siRNAs. Data is a single representative replicate from 3 independent experiments (red bar, median; dotted lines, quartiles; >5000 cells analyzed per condition). **N.** Immunoblot analysis of DOX-treated U2OS/shUb/HA-Ub(WT), U2OS/shUb/HA-Ub(K29R) and U2OS/shUb/HA-Ub(K33R) cells transfected with indicated siRNAs. **O.** Schematic model depicting regulation of SUV39H1 ubiquitylation status by TRIP12, TRABID and Cullin-RING ligases.

TRIP12 is an E3 Ub ligase containing a catalytic HECT domain required for Ub ligase activity and a WWE domain recognizing poly(ADP-ribosyl)ation (PARylation) modifications required for PAR-targeted Ub-ligase (PTUbL) activity ^26^. To validate the involvement of TRIP12 in generating nuclear K29-Ub chains, we obtained TRIP12 expression constructs encoding WT, catalytic-dead (C2007A, herein HECT*) and PAR-binding deficient (R842A, herein WWE*) mutants (**Figure 4C**), and introduced silent coding mutations providing resistance to an siRNA targeting TRIP12 (**Figure S5D**). Expression of siRNA-resistant GFP-TRIP12 restored nuclear K29-linked ubiquitylation in cells depleted of endogenous TRIP12 in a manner that was dependent on E3 ligase activity as expected, but independent of PAR-binding (**Figure 4D**; **Figure S5E**). Further suggesting a preference for TRIP12 in catalyzing K29-linked Ub chains, our unbiased MS dataset showed a clear accumulation of ubiquitylated TRIP12 along with SUV39H1, RNF168 and other proteins after Ub(K29R) replacement (**Figure 3B**). Many E3 Ub ligase enzymes have been suggested to self-regulate by auto-ubiquitylation either in *cis* or *trans* via oligomerization ^27^. Consistently, TRIP12 accumulates in Ub(K29R)-replaced cells and displays a mobility shift similar to other K29-Ub substrates (**Figure 3B**; **Figure S5F**), suggesting that TRIP12 auto-ubiquitylation via K29-Ub chains promotes its proteasomal turnover.

Given the strong evidence for TRIP12 as a major E3 ligase responsible for nuclear K29-linked ubiquitylation, we next asked whether TRIP12 is required for proteolytic turnover of SUV39H1. Depletion of endogenous TRIP12 in U2OS cells resulted in clear stabilization of SUV39H1 (**Figure 4E**), and depletion of TRIP12 removed the K29-Ub chains detected directly on SUV39H1 (**Figure 4F**). Stabilization of SUV39H1 was proteasome-dependent, as depletion of TRIP12 rescued SUV39H1 turnover in the presence of cycloheximide (**Figure 4G**), comparable to the effect of K29R-Ub replacement. Consistent with our data showing dispensable PAR-binding activity of TRIP12 for nuclear K29-Ub formation, PARylation was not required for the turnover of SUV39H1, which was degraded normally in the presence of the PARP inhibitor Olaparib (**Figure S5G**). Taken together, these results indicate that TRIP12 is the dominant writer of nuclear K29-linked Ub in human cells responsible for K29-linkage-dependent ubiquitylation and degradation of SUV39H1.

### K29-linked chains extend from K48-linked seeding and are required for efficient degradation of nuclear substrates

We next explored the role of Cullin E3 Ub ligases, depletion of which showed a modest putative effect in the maintenance of K29-Ub signal in U2OS cells (**Figure 4B**). The appearance of multiple Cullin E3 ligases (CUL1, CUL4A, CUL4B and CUL5) in our siRNA screen led us to examine the role of Cullins as a redundant grouping. To this end we utilized a potent inhibitor of Neddylation (herein NEDDi) that selectively inhibits NEDD8-activating enzyme E1 regulatory subunit (NAE1) ^28^, a core component of the UBA3-NAE1 dimer required for Cullin activation by covalent attachment of the ubiquitin-like modifier NEDD8^29^. Treatment with NEDDi impaired degradation of SUV39H1 in presence of cycloheximide, indicating a likely involvement of one or more Cullin E3 ligases in the regulation of SUV39H1 stability (**Figure 4H**). We previously observed accumulation of high-molecular weight Ub smears on K29-linked ubiquitylation substrates including SUV39H1 and RNF168 in Ub(K29R)-replaced cells that were also evident upon TRIP12 depletion (**Figure 3B Figure S5H,I**). As K29-linked Ub is known to branch from K48-linked Ub on several substrates ^22,30^, we hypothesized that TRIP12 may catalyze K29-linked extensions from preexisting K48-linked Ub chains on SUV39H1. We therefore used Ub(K29R)-replaced cells as a baseline to test if blocking the Cullin enzyme family would reduce Ub accumulation on SUV39H1. NEDDi treatment completely abolished Ub-modification on SUV39H1 in Ub(K29R)-replaced cells in a manner comparable to treatment with the E1 Ub inhibitor (**Figure 4I**; **Figure S5J**). Moreover, NEDDi treatment diminished both K48-and K29-linked Ub on SUV39H1 (**Figure S5K**). This suggests that Cullin-mediated K48-linked ubiquitylation of SUV39H1 is a prerequisite for its TRIP12-dependent K29-linked ubiquitylation. Interestingly, NEDDi treatment had no effect on Ub chains accumulating on RNF168 in K29R Ub-replaced cells (**Figure 4I**), in agreement with the well-characterized ability of the NEDDi-insensitive HECT E3 ligase UBR5 to regulate RNF168 ^23^, and the known preference for UBR5 to catalyze K48-linked Ub chains ^31^. Altogether these results suggest that TRIP12-dependent K29-linked ubiquitylation of substrates including SUV39H1 and RNF168 is primed by upstream K48-Ub generating E3 Ub ligases, the identities of which may differ between nuclear substrates.

### TRABID antagonizes K29-linked Ub chains on nuclear substrates

Ub chains are tightly regulated through the action of DUBs. Having established TRIP12 as the major writer of nuclear K29-linked Ub, we next investigated whether the proteostasis of nuclear K29-Ub substrates is regulated via DUB activity. TRABID (aka ZRANB1) is a well-characterized nuclear localizing DUB antagonizing K29- and K33-linked Ub chains ^32^, and as such we obtained a TRABID expression vector for binding studies. We introduced mutations (**Figure 4J**) to generate a catalytic-dead (C443S, herein OTU*) variant of TRABID, which has been described as a trapping mutant that binds Ub-modified substrates with increased affinity ^30^. We further generated a Ub-binding deficient variant of TRABID OTU* by introducing a T14A mutation into the NZF domain (T14A/C443S, herein NZF*) that has been shown to abolish Ub-binding of TRABID ^33^. As expected, immunoprecipitation of TRABID WT showed a weak affinity with SUV39H1, which was strongly stabilized when substituted with TRABID OTU*. The TRABID-SUV39H1 interaction was weakened again by abolishing Ub-binding with TRABID NZF* (**Figure 4K**), suggesting that TRABID recognizes Ub modifications on SUV39H1. TRABID overexpression was accompanied by a minor increase in SUV39H1 stability (**Figure S5L**). Conversely, depletion of TRABID resulted in destabilization of endogenous SUV39H1, RNF168 and NDC80 in U2OS cells (**Figure 4L**), likely due to acceleration of their K29-linked ubiquitylation in the absence of TRABID. Furthermore, the effect of TRABID depletion on SUV39H1 stability was replicated in other mammalian cell lines (**Figure S5M**). Consistently, TRABID-depleted cells showed a robust increase in nuclear K29-Ub chains when immunostained using sAB-K29 (**Figure 4M**; **Figure S5N**). Finally, destabilization of SUV39H1 in TRABID-depleted cells was rescued upon Ub(K29R) replacement (**Figure 4N**) but not Ub(K33R) replacement, indicating that TRABID activity towards SUV39H1 is K29-Ub-dependent. Taken together, these results show that SUV39H1 proteolytic turnover via Cullin-RING ligase-dependent and TRIP12-mediated K29-linked ubiquitylation is antagonized by the DUB activity of TRABID (**Figure 4O**).

### K29-Ub-dependent regulation of SUV39H1 is required for maintenance of H3K9me3 patterns

Organization of mammalian chromatin into transcriptionally repressed heterochromatin is controlled by the addition of epigenetic modifications of DNA and histones. Constitutive heterochromatin is modified by H3K9me3 ^34^, with SUV39H1 acting as the major histone methyl transferase catalyzing H3K9me3 formation in these large domains ^17^. Given the novel role of nuclear K29-Ub in regulating SUV39H1 stability, we therefore explored whether and how altered SUV39H1 stability affects the epigenetic landscape in human cells. To this end, we performed unbiased MS-based profiling of histone post-translational modifications (PTMs) in Ub(WT) and Ub(K29R) replacement cell lines treated or not with doxycycline.

Consistent with increased SUV39H1 activity, K29R-Ub replacement corresponded with a marked increase in global H3K9me3 levels (**Figure 5A**). Strikingly, however, other changes in the histone PTM landscape were negligible, apart from a minor decrease in H3K9me1 detection that most likely reflects the corresponding increase in H3K9me3 (**Figure 5A**; **Dataset S3**). We confirmed that Ub(K29R)-replaced U2OS cells do not show increased stabilization of other known H3K9me3 writers and erasers (**Figure 5B**), arguing that the increased level H3K9me3 is largely due to the stabilization of SUV39H1. Consistently, the excess SUV39H1 pool in K29R-Ub replaced cells exclusively accumulated on chromatin (**Figure S6A**). We next subjected Ub(WT) and Ub(K29R) replacement cell lines treated or not with doxycycline to quantitative ChIP-seq analysis of H3K9me3, in order to profile alterations in the H3K9me3 landscape in the absence of K29-linked Ub. Consistent with our proteomic data (**Figure 5A**), the total H3K9me3 ChIP-seq signal was increased in Ub(K29R)-replaced cells (**Figure 5C,D**). This increase in H3K9me3 levels upon loss of K29-Ub did not alter the overall H3K9me3 landscape (**Figure 5C**; **Figure S6B**). H3K9me3 domains thus remained positionally stable in Ub(K29R)-replaced cells and did not expand substantially to neighboring coding regions, despite increased H3K9me3 levels across the domains (**Figure 5C**; **Figure S6B**). These findings reveal an important role of K29-linked ubiquitylation in maintaining a proper balance of H3K9 modifications by restricting SUV39H1 levels and activity in heterochromatin.

**Figure 5.**
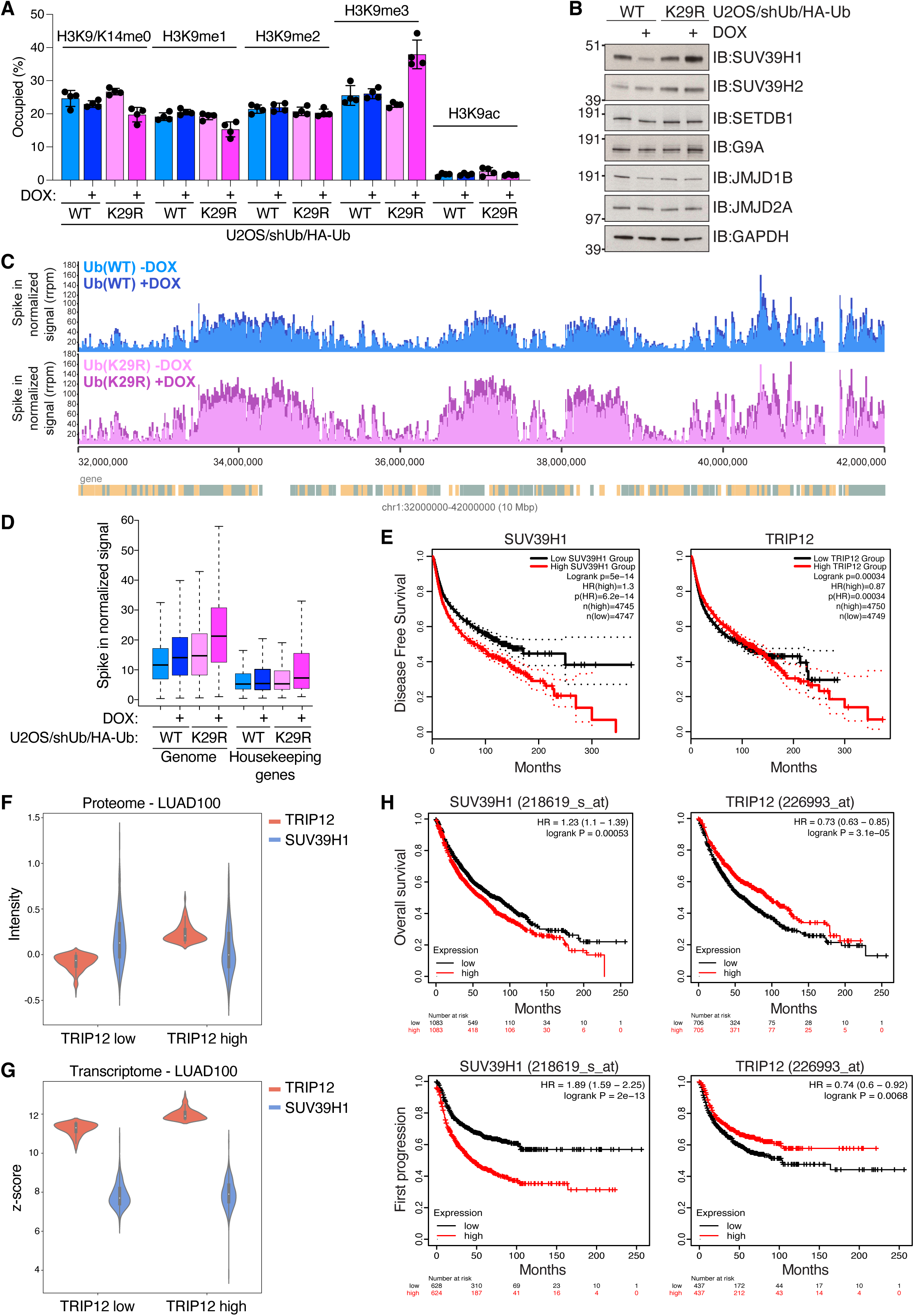
Abrogation of K29-linked ubiquitylation deregulates H3K9me3 status. **A.** Quantitative proteomic analysis of H3K9 modifications in Ub(WT) and Ub(K29R) replacement cell lines treated or not with DOX (*n*=4 biological replicates). **B.** Immunoblot analysis of Ub(WT) and Ub(K29R) replacement cell lines treated or not with DOX, using indicated antibodies. **C.** Snapshot of chromosome 1 depicting H3K9me3 ChIP-seq signal distribution in Ub(WT) and Ub(K29R) replacement cell lines treated or not with DOX. **D.** ChIP-Seq quantification of H3K9me3 in Ub(WT) and Ub(K29R) replacement cell lines treated or not with DOX stratified by genomic location. Reads per million normalized to *Drosophila* spike-in (*n*=4 biological replicates). **E.** Kaplan-Meier analysis of disease-free survival stratified by expression of SUV39H1 (left) or TRIP12 (right) in a pan-cancer cohort. Dotted lines represent 95% confidence interval. Analysis was performed using GEPIA2 on 33 cancer types. Hypothesis evaluation was performed with log-rank test. Hazard ratios were calculated according to the Cox proportional-hazards model. **F.** LUAD100 proteome levels of indicated proteins stratified by relative high and low TRIP12-expressing samples. Data are shown as log2 ratio of reporter ion intensity. **G.** LUAD100 transcriptome levels stratified by relatively high and low TRIP12-expressing samples. **H.** Kaplan-Meier analysis in non-small cell lung cancer (NSCLC) stratified on SUV39H1 (left) or TRIP12 (right) expression. Overall survival (upper) and first progression-free survival (lower panel) are shown. Analysis was performed using KMPlotter tool. Hazard ratios and p-values were calculated through Cox regression.

### TRIP12 and SUV39H1 expression levels anticorrelate in cancer

Heterochromatin establishment and maintenance is central to cell identity. Aberrant H3K9me3 occupancy is associated with loss of cell identity, such as premature aging or cancer ^35^. As such we next explored whether TRIP12-dependent regulation of SUV39H1 is disrupted in a disease context. Kaplan-Meier analysis of survival across a dataset of 33 types of cancer (GEPIA2 ^36^) showed that higher expression of SUV39H1 correlates with poorer disease-free survival (**Figure 5E**). Conversely, the same cohort showed the opposite relationship for TRIP12, with poorer disease-free survival correlating with lower TRIP12 expression (**Figure 5E**). Although higher expression at the mRNA level is likely independent of Ub-mediated degradation, poorer disease-free survival with increased SUV39H1 may indicate a generally adverse effect of high SUV39H1 in cancer progression, highlighting the importance of regulating SUV39H1 abundance and activity.

We next explored the relationship between TRIP12 and SUV39H1 protein levels in various cancer types. Lung adenocarcinoma (LUAD) is one of the most common subtypes of non-small cell lung cancer (NSCLC) which makes up 85% of cases worldwide ^37^. Stratification of LUAD samples on TRIP12 protein detection showed lower SUV39H1 protein detection in “high” TRIP12 expressing samples (**Figure 5F**), a relationship not seen when samples were stratified on TRIP12 mRNA levels (**Figure 5G**). Consistently, Kaplan-Meier analysis of survival in a cohort of NSCLC patients ^38^ showed a favorable outcome for patients with either lower SUV39H1 or high TRIP12 expression in both overall survival and first progression, emphasizing the strong anticorrelation of these proteins in survival (**Figure 5H**).

## Discussion

Low abundance Ub chains are notoriously difficult to isolate for study due to a paucity in high performance linkage-specific Ub detection reagents. To remedy this barrier, we have leveraged a robust cell-based strategy for the conditional abrogation of all seven individual lysine-based Ub-chain topologies in human cells, expanding our previous work identifying a role for K27-linked Ub chains in p97-dependent nuclear proteostasis ^16^, and have carefully benchmarked performance using biochemical and cell biological validation. The Ub replacement cell line panel described here complements an array of tools to study the ubiquitin-proteasome system at multiple levels, such as Absolute Quantitation of Ubiquitin linkage Abundance (Ub-AQUA) ^39^ and Ubiquitin Chain Restriction (UbiCRest) analysis ^40^ for the respective analysis of Ub chain composition and architecture, the DegronID database^41^ for mapping of E3 Ub ligase specificity, Ubiquitin interactor Affinity enrichment-MS (UbiA-MS) ^10^ for the profiling of Ub chain interacting proteins, and a proteome half-life resource ^25^. Our Ub replacement cell lines were rigorously selected to ensure uniform conditional expression of exogenous Ub at levels similar to that of endogenous Ub and thus maximal compliance with endogenous ubiquitylation dynamics and cell fitness upon Ub replacement. This cell line panel represents a readily available and highly accessible system for the investigation of low abundance Ub chains and their cellular functions in the absence of appropriate enrichment reagents, which often display cross reactivity with more abundant Ub-linkages, or sub-optimal performance *in vivo* as we show for a range of commercially available linkage-specific Ub affinity reagents. The system is maintained in standard cell culture conditions, requiring minimal specialized equipment or expertise to implement.

A major advantage of leveraging the Ub replacement strategy is the retention of cellular context, both in terms of abrogating a specified Ub-linkage in live cells and in keeping other Ub-substrate linkages intact. Compatibility of the Ub replacement system with live cell assays allowed us to explore the requirement of low abundance Ub-linkages for cell viability, showing a dispensability for K6-, K11-, K29- and K33-linked Ub for short-term viability of non-stressed U2OS cells. However it is likely that increased toxicity would be observed in these cell lines under appropriate perturbation conditions. K6-linked ubiquitylation, for example, was recently shown to promote the resolution of RNA-protein crosslinks (RPCs) ^8,9^. Indeed, using Ub(K6R) replacement cells we validated a strong RPC-dependent increase in K6-linked ubiquitylation. In general, our panel of Ub replacement cell lines enables rapid and efficient probing of the requirement of specific Ub-linkages for cellular processes of interest. Illustrating this, we show that K63- and K27-linked Ub chains, but not other Ub linkages, are required for Ub-dependent recruitment of 53BP1 to damaged DNA, in line with previous findings ^12,13^.

A main utility of the Ub replacement strategy lies in its unique ability to inform on how disrupting the formation of specific Ub-linkages impacts cellular ubiquitylation processes globally. This offers insights into the scope of proteins impacted by a particular Ub-linkage type and thus the cellular functions of this Ub chain topology. In the present study, we provide system-wide proteomic inventories of ubiquitylome and whole proteome changes induced by abrogating each lysine-based Ub chain type under otherwise unperturbed conditions in human U2OS cells, providing a valuable resource to further explore cellular roles of linkage-specific ubiquitylation processes. For instance, this revealed prospective roles of K33- and K27-linked ubiquitylation in regulating RNA processing and mitochondrial proteins, respectively. Moreover, we discovered a notable enrichment of nuclear and chromatin-associated factors among ubiquitylated proteins responsive to disruption of K29-linked ubiquitylation, suggesting a key role of K29-Ub in regulating chromatin status and dynamics. Indeed, we demonstrate that SUV39H1, a major writer of the repressive histone mark H3K9me3, is a prominent target of regulation by K29-linked Ub chains, which we found are instrumental for proteasomal degradation of SUV39H1. In fact, the Ub replacement system proved particularly useful for identification of targets of K29-linked ubiquitylation, due to a striking tendency of substrates to accumulate K48-linked Ub in the absence of K29-Ub. This behavior indicates that, at least for some K29-Ub targets such as SUV39H1, upstream K48-linked ubiquitylation was insufficient for degradation in line with recent findings ^22^. K29-linked Ub chains were recently shown to act in p97-dependent degradation ^6^, and it is conceivable that the presence of K29-linked Ub promotes substrate unfolding required prior to degradation initiation at the proteasome, e.g. by facilitating extraction of substrates tightly bound to chromatin or other macromolecular structures where K48-Ub signals alone are ineffective. We identified TRIP12 and TRABID as the major writer and eraser of nuclear K29-linked Ub modifications, respectively, and we establish that these enzymes are the key effectors controlling K29-linked ubiquitylation of SUV39H1. We also show that TRIP12 activity towards RNF168 is K29-Ub-dependent, consistent with a previously identified role TRIP12 in the regulation of the nuclear RNF168 pool ^23^.

Modulation of substrate stability by the coupling of K48- and K29-linked Ub signals showed a clear directionality, with K48-linked Ub addition occurring upstream of TRIP12 activity, catalyzed by other E3 Ub ligases whose identity may differ between individual K29-Ub substrates as we demonstrate for SUV39H1 and RNF168. We found that the K48-Ub modifications on SUV39H1 priming its K29-linked ubiquitylation were fully dependent on Cullin-RING E3 ligase activity, consistent with previous studies implicating Cullin-RING complex components in SUV39H1 degradation ^42,43^. Depletion of nuclear-localizing TRIP12 left cytoplasmic K29-Ub levels largely unchanged, indicating a compartmentalization of K29-Ub activity and non-nuclear K29-Ub writers whose identities remain to be established.

Histone modifications are a central component of epigenetic control of gene expression, however how epigenetic writers are themselves controlled is an area that remains largely understudied. We identified an important role for K29-linked Ub chains in the maintenance of the cellular epigenetic landscape that selectively impinges on H3K9 methylation status. Contrary to the impact on SUV39H1, we found no evidence of altered abundance of a range of other known H3K9me3 writers and erasers with K29-Ub abrogation, suggesting that the role of this Ub linkage type in regulating H3K9me3 status may be predominantly exerted via TRIP12-dependent stimulation of SUV39H1 turnover. Notably, while disrupting K29-Ub-dependent degradation of SUV39H1 significantly increased total H3K9me3 abundance, this predominantly manifested as hyper-accumulation of this mark within existing H3K9me3 domains, indicative of deregulation of enzyme activity rather than targeting mechanism. Why SUV39H1 is uniquely controlled by K29-linked ubiquitylation among H3K9me3 regulatory factors remains to be addressed. During cell division, H3K9me3 is established with slow kinetics on new histones, limiting accumulation of this repressive modification ^44^. Tight control of SUV39H1 levels might be needed to control H3K9me3 propagation, matching maintenance via SUV39H1 read-write function ^34,45^ with cell cycle speed. It will be interesting to explore whether K29-Ub-mediated control of SUV39H1 plays a role in reorganization of H3K9me3 patterns during development and/or erasure of H3K9me3 during cellular reprogramming. Aberrant epigenetic regulation is a prevalent feature of multiple cancer types ^46^. We show here a putative disparity in survival outcomes for cancer patients when stratified by expression of both TRIP12 and SUV39H1. This was particularly evident for NSCLC patients, a subgroup of which also showed lower SUV39H1 protein levels in cancer samples with relatively higher TRIP12 that was associated with a favorable prognosis. These observations emphasize the strong anti-correlative relationship between SUV39H1 and TRIP12 and its potential importance for fitness and survival.

In summary, we describe here a robust panel of cell lines for conditional abrogation of lysine-based Ub topologies with broad applicability across cell biology, exemplified by the discovery of a novel role for K29-linked Ub in promoting epigenome maintenance via regulation of SUV39H1 stability, with implications for cell physiology and disease.

## Methods

### Plasmids and siRNAs

Plasmids encoding Ub depletion (pPuro-shUb) and Ub replacement (pcDNA5/FRT/TO-HA-Ub WT, K27R, K29R, K33R and K63R) were described previously ^15,16^. In brief, shUb encodes siRNA sequences 5’-ACACCATTGAGAATGTCAA-3’ targeting *UBC* and *UBA52*, and 5’-AGGCCAAGATCCAGGATAA-3’ targeting *UBB* and *RPS27A*. Ub replacement casettes to achieve K6R, K11R and K48R replacement were generated via sequential mutagenesis of pcDNA5/FRT/TO-HA-Ub(WT). All mutagenesis was performed using the Q5 Site-Directed Mutagenesis kit (New England Biolabs) and the following primer sets: UBA52 K6R: 5’-ATC TTT GTG AGG ACC CTC ACT GG-3’ and 5’-CTG CAT GGT GTA CCA GCC-3’; RPS27A K6R: 5’-ATT TTC GTG AGA ACC CTT ACG-3’ and 5’-CTG CAT AGC GTA ATC TGG-3’; UBA52 K11R: 5’-CTC ACT GGC AGA ACC ATC ACC-3’ and 5’-GGT CTT CAC AAA GAT CTG C-3’; RPS27A K11R: 5’-CTT ACG GGG AGG ACC ATC ACC-3’ and 5’-GGT TTT CAC GAA AAT CTG CAT AG-3’; UBA52 K48R: 5’-TTT GCC GGC AGA CAG CTG GAG-3’ and 5’-TAT CAG ACG CTG CTG GTC-3’; RPS27A K48R: 5’-TTT GCT GGC AGG CAG CTG GAA-3’ and 5’-GAT CAG TCT CTG CTG ATC AGG-3’.

A synthetic cDNA sequence (Thermo) encoding the MultiDsk ^47^ tandem ubiquitin binding entity (TUBE) was cloned into pFN18A-HaloTag (Promega) to create pFN18A-HaloTag-MultiDsk-6xHis. GFP-SUV39H1 and FLAG-SUV39H1 were generated by Gateway recombination between pENTR221-SUV39H1 (Invitrogen Ultimate ORF IOH6289) and pDEST-53 (Thermo) and pDEST/FRT/N1-FLAG respectively (Thermo). HA-TRABID was generated by cloning TRABID cDNA into pHM6 vector (Roche). Catalytic-dead HA-TRABID C443S as per ^30^ was generated from HA-TRABID using the primer set: 5’-GCA GGA GAC TCCC TAC TTG ATT C-3’ and 5’-AGT CCG GTT CCA AAG TGC-3’. Ub-binding deficient HA-TRABID T14A/C443S as per ^33^ was generated from HA-TRABID C443S using the primer set: 5’-TGA ATA TTG TGC GTA TGA AAA CTG-3’ and 5’-CAA GCC CAC TTA ATT CCA C-3’. GFP-TRIP12 and variants encoding HECT* and WWE* (C2034A and R869A respectively) were a kind gift from Matthias Altmeyer ^26^ and were altered with silent mutations for siRNA resistance using the following primer set: 5’-ACAGTTTTAGAGATGGATTTGAATCAG -3’ and 5’-CGAACTGCCTAGAAACGCCTTC -3’.

The following siRNAs from Sigma were used: siMM1 (5′-GGGAUACCUAGACGUUCUA-3′); siTRIP12-1 (5’-GCAAUUUGAUUCGUUCAGA-3’); siTRIP12-2 (5’-GCACCUAGAUUGGAUAGAA-3’); siTRIP12-4 (5’-AAAGUAAUAAAGAUUGUGU-3’); siTRIP12-5 (5’-GCAAUUUGAUUCGUUCAGA-3’); siTRABID-1 (5’-GAAUCGUCCUUCUGCCUUU-3’); siTRABID-2 (5’-GUGAUCAUCCCAGACCUAA-3’); siSUV39H1-1 (5’-AGAACAGCUUCGUCAUGGA-3’); siSUV39H1-2 (5’-GGCCUUCGUGUACAUCAAU-3’); siSUV39H1-3 (5’-CAAAUCGUGUGGUACAGAA-3’). All siRNAs were double-transfected at 20 nM using Lipofectamine RNAiMAX reagent (Thermo) according to manufacturer’s instructions. All siRNA transfections were harvested at 72 h post initial transfection unless otherwise indicated.

E3 ligase screening was performed using the Human On-Target Ubiquitin Conjugation siRNA libraries subset 1 (DHARG-105615, Horizon), subset 2 (DHARG-105625, Horizon) and Subset 3 (DHARG-105635, Horizon). Cells were plated in 96-well black-walled optical imaging culture vessels 24 h prior to double-transfection at 50 nM, fixed after 72 h before staining as indicated and high content image acquisition (see immunofluorescence). Sample mean intensities were normalized to siMM1 within each 96-well plate. Z-score was calculated as:

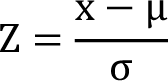

where x is sample intensity, µ is population mean and σ is the population standard deviation.

### Cell culture

Human cell lines (U2OS, HeLa and RPE1) were obtained from ATCC, grown under standard conditions at 37 °C and 5% CO_2_ in DMEM (Thermo) supplemented with 10% FBS (v/v) and Penicillin-Streptomycin (Thermo). TIG3 cells were previously described ^44^. All Ub replacement cell lines were maintained under selection of Puromycin (Sigma), Blasticidin and Hygromycin B (Thermo). Stable clones were carefully validation for uniform Ub expression. Where indicated, Ub replacement was achieved by doxycycline (DOX) (0.5 µg/ml) treatment. All cells were cultured in humidified incubators containing 5% CO_2_ at 37°C and were regularly tested negative for Mycoplasma infection. The cell lines were not authenticated.

U2OS cells stably expressing GFP-RNF168 (U2OS/GFP-RNF168) were described previously ^48^. U2OS/GFP-SUV39H1 cells were generated by transfection and selection with G418 (Invivogen) at 400 µg/ml. Drosophila S2-DRSC suspension cells were cultivated at 25°C with 5% CO_2_ in spinners using M3+BPYE media: comprising Shields and Sang M3 Insect Medium (Sigma), KHCO3 (Sigma), yeast extract (Sigma), bactopeptone (BD Biosciences), 10% heat-inactivated FBS (GE Hyclone), and 1X penicillin/streptomycin (Gibco). hTERT TIG3 fibroblasts from fetal lung were additionally supplemented in culture with 10% non-essential amino acids.

Incucyte confluence assays were performed as described ^16^. In brief, cellular confluence was assessed using the Incucyte S3 Live-Cell Analysis System (Sartorius). Unless otherwise indicated, the following standard treatment concentrations were used: doxycycline (1µg/ml, Sigma), MLN-7243 (E1i; 5 μM, Active Biochem), NMS-873 (5 μM, Sigma), MG132 (20 μM; Sigma), Actinomycin D (ActD; 2 μg/ml, Sigma), MLN-4924 (NEDDi; 1 µM, Selleckchem), Cycloheximide (CHX; 20 µg/ml, Sigma), MZ1 (500 nM, R&D Systems) and Olaparib (1 µM, AstraZeneca). DNA double-strand breaks (DSBs) were induced where indicated using the XYLON Smart X-ray system.

### RT-qPCR

Total RNA was extracted from cells following the manufacturer’s protocol (RNeasy kit, Qiagen). cDNA was synthesized from RNA by reverse transcription PCR (iScript cDNA Synthesis Kit, Bio-Rad). Real-time quantitative PCR was performed using the Stratagene Mx3005P System and Brilliant III Ultra-Fast SYBR Green QPCR Master Mix (Agilent). GAPDH mRNA levels were used as control for normalization. The following primers were used for amplification of the respective cDNAs: GAPDH (forward): 5′-CAGAACATCATCCCTGCCTCTAC-3′; GADPH (reverse): 5′-TTGAAGTCAGAGGAGACCACCTG-3′; HA-RPS27A (forward): 5′-TACCCTTACGATGTACCAGA-3′; HA-RPS27A (reverse): 5′-GAGGGTTCAACCTCGAGGGT-3′; RPS27A (forward): 5′-TCGTGGTGGTGCTAAGAAAAGG-3′; RPS27A (reverse): 5′-TTCAGGACAGCCAGCTTAACCT-3; UBB (forward): 5′-CTTTGTTGGGTGAGCTTGTTTGT-3′; UBB (reverse): 5′-GACCTGTTAGCGGATACCAGGAT-3′; UBA52 (forward): 5′-CTGCGAGGTGGCATTATTGAG-3′; UBA52 (reverse): 5′-GTTGACAGCACGAGGGTGAAG-3′; UBC (forward): 5′-GTGTCTAAGTTTCCCCTTTTAAGG-3′; UBC (reverse): 5′-TTGGGAATGCAACAACTTTATTG-3′.

### In silico local deformation prediction of single amino acid mutation

The amino acid sequence for WT and single K-to-R mutants of human Ub (P0CG48, Uniprot) were input to AlphaFold2_mmseqs2 via ColabFoldv1.5.5 ^49^ with standard settings and no template mode. Output structures were input to PDAnalysis ^50^. Effective strain between the WT structure and individual mutations were calculated using an average of 5 output structures from AlphaFold2. Effective strain was then superimposed onto the default structure using the b-factor column via a custom Python3 script. All protein structures were visualized with ChimeraX v1.7.

### Antibodies

The following primary antibodies were used: Actin (MAB1501, Millipore (RRID:AB_2223041)); GAPDH (sc-20357, Santa Cruz (RRID:AB_641107)); GFP (11814460001, Merck (RRID:AB_390913)); HA (11867423001, Roche (RRID;AB_390918)), FLAG (F1804, Sigma (RRID:AB_262044)); RPS27A (ab111598, Abcam (RRID:AB_10863285)); Tubulin (T9026, Sigma (RRID:AB_2541185)); UBA52 (PA5-23685, Thermo (RRID:AB_2541185)); Ub (FK2) (BML-PW8810, Enzo (RRID:AB_2820835)); Ub (P4D1) (sc-8017, Santa Cruz (RRID:AB_2762364)); Ub K11-linked (SAB5701121, Sigma); Ub K27-linked (ab181537, Abcam (RRID:AB_2713902)); Ub K33-linked (PA5-120623, Thermo (RRID:AB_2914195)); Ub K48-linked (Apu2) (05-1307, Sigma (RRID:AB_11213655)); Ub K63-linked (BML-PW0600-0025, Enzo (RRID:AB_2052278)); SUV39H1 (8729T, Cell Signaling (RRID:AB_10829612)); SUV39H2 (LS-C116360, Ls Bio (RRID:AB_10914491)); TRABID (ab262879, Abcam); TRIP12 (A301-814A, Bethyl Labs (RRID:AB_1264344)); JMJD1B (2621S, CST (RRID:AB_915946)); JMJD2A (NB110-40585, Novus (RRID:AB_669535)); RNF168 (kind gift from Daniel Durocher ^51^); NDC80 (kind gift from Jakob Nilsson; raised against NDC80 aa1-230 by Moravian Biotechnology); H3K9me3 (ab176916, Abcam (RRID:AB_2797591)); SETDB1 (ab107225, Abcam (RRID:AB_10861045)); BRD4 (13440, CST (RRID:AB_2687578)); Histone H3 pS10 (06-570, Merck (RRID:AB_310177)); 53BP1 (MAB3802, Millipore (RRID:AB_11212586)); ψH2AX (05-636, Millipore (RRID:AB_309864)).

### Cell lysis, immunoblotting and immunoprecipitation

Immunoprecipitation (IP) of HA-tagged proteins was performed using Anti-HA Affinity matrix (1181501600, Roche) under denaturing conditions as described ^16^. Cell lysis was performed using RIPA buffer (140 mM NaCl; 10 mM Tris-HCl (pH 8.0); 0.1% sodium deoxycholate (w/v); 1% Triton X-100 (v/v); 0.1% SDS (w/v); 1 mM EDTA; 0.5 mM EGTA). GFP-trap and DYKDDDDK Fab-Trap (Chromotek) pulldown was performed according to manufacturer’s instructions. Unless indicated all whole lysate and immunoprecipitation samples were prepared and washed in non-denaturing buffer (150 mM NaCl; 50 mM Tris-HCl, pH 8.0; 0.5mM EDTA). All samples obtained under denaturing conditions were lysed and washed in denaturing buffer (50 mM NaCl; 20mM Tris-HCl, pH 7.5; 0.5% sodium deoxycholate (w/v); 0.5% igepal (v/v); 0.5% SDS (w/v); 1 mM EDTA). All samples indicated as subject to MultiDsk pulldown were lysed and washed in a stringent buffer (50 mM Tris-HCl, pH 8.0; 1 M NaCl; 1% Igepal v/v; 0.1% SDS w/v). Halo-MultiDsk was conjugated to HaloLink Resin (Promega) prior to IP for 1 h at RT in wash buffer (100 mM Tris-HCl, pH 7.5; 150 mM NaCl; 0.05% Igepal v/v). All lysis and wash buffers were supplemented with PMSF Protease Inhibitor (1 mM, Thermo), EDTA-free protease inhibitor Cocktail Tablets (Roche), N-ethylmaleimide (1.25 mM, Sigma) and DUB inhibitor PR-619 (50 µM, Calbiochem).

Met-Gly-Ser-His6-3C-codon-optimized LotA-N (catalytic inactive C13A mutant) with a C-terminal (Ala-Ser)3 linker followed by an AviTag (Avidity) was expressed from a pET vector in *E. coli* (ArcticExpress (DE3), Agilent), and purified as previously described ^8^. Following cell lysis in modified RIPA buffer, pre-cleared protein lysate was incubated with 1 μM biotin-conjugated LotA-N for 2 h at 4 °C. High capacity Neutravidin beads (15 μl per sample) were added and incubated for 1 h at 4 °C on a rotating wheel to precipitate biotin-conjugated probes. Beads were washed four times with modified RIPA buffer supplemented with protease inhibitors and 10 mM N-ethylmaleimide to remove nonspecific binders prior to elution in NuPAGE LDS 2×Sample Buffer (Life Technologies).

### Chromatin fractionation

U2OS cells were collected by scraping in ice-cold PBS and subsequent centrifugation for 5 min at 400g. The pellets were lysed with ice-cold cell lysis buffer (10 mM Tris, pH 8.0; 10 mM KCl; 1.5 mM MgCl_2_; 0.34 M sucrose; 10% glycerol; 0.1% Triton X-100) supplemented with protease, phosphatase and DUB inhibitors. After centrifugation for 5 min at 2000g, the soluble fraction was recovered. The pellets were carefully washed once in lysis buffer and centrifuged as in the previous step. Pellets were resuspended in RIPA buffer supplemented with benzonase and protease, phosphatase and DUB inhibitors and incubated in a thermomixer for 15 min at 37 °C. A last centrifugation step (16000g for 10 min at 4 °C) allowed the recovery of the solubilized chromatin-bound proteins.

### Immunofluorescence

Coverslips were fixed in 10% formalin buffer (VWR) for 15 min at room temperature, permeabilized with 0.5% Triton-X for 5 min and blocked with 5% BSA (Sigma). Cells were stained with primary antibody for 1 h at RT, washed with PBS and stained with a combination of Alexa Fluor secondary antibodies (Thermo) and 4’,6-Diamidino-2-Phenylindole (DAPI; Molecular Probes) for 30 min at RT. Coverslips were then washed, dried and mounted using Mowiol (Sigma). Where indicated, cells were pre-extracted prior to fixation using CSK buffer (100 mM NaCl; 10 mM HEPES; 3 mM MgCl_2_; 300 mM sucrose; 0.25% Triton-X; 1 mM PMSF). Where indicated, nascent DNA synthesis was estimated by incubation with 10 μΜ 5-ethynyl-2’-deoxyuridine (EdU; Thermo) for 1 h prior to fixation.

EdU incorporation was labelled using Click-iT Plus EdU Alexa Fluor 647 Imaging Kit (Thermo) according to manufacturer’s instructions. Transcription and translation efficiency was assessed as described ^16^. Quantitative image-based cytometry (QIBC) was performed as described ^16^. In brief, images were acquired and analyzed using the ScanR high-content screening platform (Olympus). Cell cycle gating was performed using DAPI and EdU staining, with mitotic figures determined as a percentage of whole population based on histone H3-pS10 staining. Representative images were acquired with a confocal microscope (LSM 880; Carl Zeiss), mounted on a confocal laser-scanning microscope (Zeiss AxioObserver.Z1; Carl Zeiss) equipped with a Plan-Apochromat 40×/1.3 NA oil immersion objective. Image acquisition was performed with ZEN 2.1 software (Carl Zeiss). Raw images were exported as TIFF files and identical settings were used on all images of a given experiment when adjustments in image brightness were applied.

### Mass spectrometry (whole proteome analysis)

Protein samples were collected in modified RIPA buffer (50 mM Tris, pH 7.5; 150 mM NaCl; 1 mM EDTA; 1% NP-40; 0.1% sodium deoxycholate) supplemented with protease inhibitors (Complete Protease Inhibitor Cocktail Tablets, Roche), 1 mM sodium orthovanadate, 5 mM β-glycerophosphate, 5 mM sodium fluoride and 10 mM N-ethylmaleimide. Chromatin-bound proteins were extracted by addition of NaCl to a final concentration of 500 mM and pulse sonication at 4 °C for 10 min. Total protein concentrations were estimated using the QuickStart Bradford Protein assay (Bio-Rad) from the pre-cleared lysates. Proteins were precipitated in fourfold excess of ice-cold acetone and subsequently re-dissolved in denaturation buffer (6 M urea; 2 M thiourea; 10 mM HEPES, pH 8.0). Cysteines were reduced with 1 mM dithiothreitol and alkylated with 5.5 mM chloroacetamide. Proteins were digested with MS-approved Trypsin (Sigma). Protease digestion was stopped by addition of trifluoroacetic acid to 0.5%, and precipitates were removed by centrifugation. Peptides were purified and desalted using reversed-phase Sep-Pak C18 cartridges (Waters) and eluted in 50% acetonitrile. For full proteome profiling the peptides were supplemented with trifluoroacetic acid up to 1% and analysed on a quadrupole Orbitrap mass spectrometer (Astral, Thermo Scientific) equipped with a UHPLC system (Vanquish, Thermo Scientific). Raw data files were converted to mzML format using MSConvert and analyzed using FragPipe (development v.21.1) LFQ-MBR workflow ^52–54^. Parent ion and MS2 spectra were searched against a database containing 98,566 human protein sequences obtained from UniProtKB (April 2018 release) using the MSFragger search engine. Spectra were searched with a mass tolerance of 6 ppm in MS mode, 20 ppm in HCD MS2 mode and strict trypsin specificity, allowing up to two miscleavages. Cysteine carbamidomethylation was searched as a fixed modification. Peptide probability was determined by the highest supporting PSM probability. Results for each experiment were aggregated and filtered at 1% peptide FDR. Data analysis and visualization were performed using RStudio in the R 4.4.1 environment.

### Mass spectrometry (HA-tag affinity purification)

HA-tag affinity purification was performed using Anti-HA Affinity Matrix (Roche). Partial on-bead digest was performed with buffer (2 M urea; 2 mM DTT; 20 µg/ml Trypsin; 50 mM Tris, pH 7.5) incubated at 37 °C for 30 min at 1,400 rpm. Supernatants were transferred to new tubes, alkylated with 25 mM chloroacetamide (CAA) and further digested overnight at RT and 1,000 rpm. Digestion was terminated with 1% trifluoroacetic acid (TFA). Peptides were desalted and purified using styrenedivinylbenzene-reversed phase sulfonate (SDB-RPS) StageTips prepared in 0.2% TFA. Peptides were washed and eluted with elution buffer (80% acetonitrile (ACN); 1% ammonia) prior to vacuum-drying. Dried peptides were reconstituted in 2% ACN and 0.1% TFA.

All samples were loaded onto Evotips Pure and measured with a data-independent acquisition (DIA) method. 200 ng of peptides were partially eluted from Evotips with <35% acetonitrile and analyzed with an Evosep One LC system (Evosep Biosystems) coupled online to an Orbitrap mass spectrometer (Orbitrap Astral, Thermo Fisher Scientific) ^55–57^. Eluted peptides were separated on a 8-cm-long PepSep column (150 µm inner diameter packed with 1.5 μm of Reprosil-Pur C18 beads (Dr Maisch)) in a standard preset gradient method (21 min, 60 samples per day) with a stainless emitter (30 µm inner diameter). The mobile phases were 0.1% formic acid in liquid chromatography (LC)–MS-grade water (buffer A) and 0.1% formic acid in acetonitrile (buffer B). Data were acquired in DIA mode. Each acquisition cycle consisted of a survey scan at a resolution of 240,000 (normalized automatic gain control target (AGC) of 500% and a maximum injection time of 100 ms. Fragment ion scans were recorded with a maximum injection time of 5 ms and with 200 windows of 3Th scanning from 380 − 980 m/z. Higher-energy collisional dissociation (HCD) fragmentation was set to a normalized collision energy of 25%.

Raw files were analyzed with directDIA workflow in Spectronaut v.18.6 ^58^ using Default settings. Data filtering was set to ‘Qvalue’. ‘Cross run normalization’ was enabled with the strategy of ‘local normalization’ based on rows with ‘Qvalue complete’. FDR was set to 1% at both the protein and peptide precursor levels. Raw data was searched against the human proteome reference database, including isoform information (Uniprot March 2023). Data was filtered for 60% valid values across WT or Mutant samples (protein groups with >40% missing values were excluded from downstream statistical analysis). Protein intensities were then log2 transformed for downstream statistical and bioinformatics analysis. Imputation of missing data was performed by random numbers drawn from a normal distribution with a width of 0.3 and downshift of 1.8 applied to each sample. Significance was determined through unpaired two-sided t-test corrected by Benjamini–Hochberg at FDR of 0.1 and fold change of 1.5 ^59,60^. GO-term enrichment was performed using ClusterProfiler ^61^. Data analysis and visualization were performed using Jupyter Notebook (Python 3.9) and RStudio (v2023.03.1+446) with R (v4.3.0) for ClusterProfiler-based analysis.

### Ubiquitin-modified proteome (diGly) analysis

For the analysis of ubiquitin-modified proteomes, protein samples were collected in modified RIPA buffer from cells seeded in 150 mm cell culture dishes. Chromatin-bound proteins were extracted by the addition of NaCl to a final concentration of 500 mM and pulse sonication at 4 °C for 10 min. Total protein concentrations were estimated using the QuickStart Bradford Protein assay (Bio-Rad) from the pre-cleared lysates and a minimum of 1 mg of protein from each samples was subjected to OtUBD pulldown ^62^. Typically, 15 μl of OtUBD resin (bed volume) was used for each 1 mg of lysate protein and the mixture was incubated at 4 °C for 2 hours with rotation. The resin was allowed to settle by gentle centrifugation at 1000g x 1 min. The unbound solution was drained and the pelleted OtUBD resin was washed three times with 8 M Urea and one time with 1% SDS-PBS prior to elution. Bound proteins were eluted by incubating the resin with 2x resin volumes of 2x LDS sample buffer for 15 min at RT with rotation.

The eluted Ub-modified proteins were reduced using 5 mM DTT and then alkylated with the addition of 11 mM chloroacetamide in the dark at room temperature for 30 min. Samples were subsequently cleaned up by SP3 and digested in solution using Trypsin at 37 °C with trypsin for 16 h. Protease:protein ratios were estimated at 1:50. Digested peptides were desalted on reverse-phase C18 StageTips prior to MS analysis.

Raw data files were analyzed using MaxQuant (development version 1.5.2.8). Parent ion and MS2 spectra were searched against a database containing 98,566 human protein sequences obtained from UniProtKB (April 2018 release) using the Andromeda search engine. Spectra were searched with a mass tolerance of 6 ppm in MS mode, 20 ppm in HCD MS2 mode and strict trypsin specificity, allowing up to three miscleavages. Cysteine carbamidomethylation was searched as a fixed modification, whereas protein N-terminal acetylation and methionine oxidation were searched as variable modifications. The dataset was filtered based on posterior error probability (PEP) to arrive at a false discovery rate (FDR) of less than 1% estimated using a target-decoy approach.

### Mass spectrometry analysis of histone PTMs (EpiQMAx)

U2OS/shUb/HA-Ub(WT) and U2OS/shUb/HA-Ub(K29R) cells were seeded in quadruplicate 15-cm dishes at a density of 1 million cells per dishes and treated or not with DOX for 72 h. Cells were trypsinized and washed twice with PBS prior to collection in media. From this suspension, 10 million cells were pelleted at 300g for 5 min and washed twice with PBS. The cell pellets were snap-frozen and shipped on dry ice to EpiQMAx GmbH for further processing. Acid-extracted histones were resuspended in Laemmli sample buffer, separated by a 14–20% gradient SDS–PAGE, and stained with Coomassie (Brilliant Blue G-250, 35081.01). Protein bands within the 15–23 kDa molecular weight range were excised as single bands, destained in a 50% acetonitrile/50 mM ammonium bicarbonate solution, and chemically modified by propionylation for 30 min at room temperature with 2.5% propionic anhydride (Sigma-Aldrich) in ammonium bicarbonate, pH 7.5. Proteins were then digested overnight with 200 ng of trypsin (Promega) in 50 mM ammonium bicarbonate. The resulting supernatant was desalted using C18-Stagetips (reversed-phase resin) and carbon Top-Tips (Glygen) according to the manufacturer’s instructions, speed vacuumed until dry, and stored at -20 °C until mass spectrometry analysis.

Peptides were injected in an Ultimate 3000 RSLCnano system (Thermo-Fisher Scientific, San Jose, CA) and separated on a 15-cm analytical column (75 μm ID with ReproSil-Pur C18-AQ 2.4 μm (Dr. Maisch)) using a gradient from 4% B to 90% B (solvent A 0.1% FA in water, solvent B 80% ACN, 0.1% FA in water) over 90 min at a flow rate of 300 nl/min. The effluent from the HPLC was directly electro-sprayed into a Exploris 240 mass spectrometer (Thermo Fisher Scientific, San Jose, CA). The mass spectrometer was operated in data-dependent mode to automatically switch between full scan MS and MS/MS acquisition.

Survey full scan MS spectra (from m/z 375 to 1,600) were acquired with resolution R = 60,000 at m/z 400 (AGC target of 3 × 106). The 10 most intense peptide ions with charge states between 2 and 5 were sequentially isolated to a target value of 1 × 105 and fragmented at 27% normalized collision energy. Typical mass spectrometric conditions were as follows: spray voltage, 1.5 kV; no sheath and auxiliary gas flow; heated capillary temperature, 250 °C; and ion selection threshold, 33,000 counts.

Raw files were analyzed using Skyline software against histone H3 and H4 peptides and their respective PTMs, with a precursor mass tolerance of 5 ppm. The chromatogram boundaries of +2 and +3 charged peaks were validated, and the Total Area MS1 under the first four isotopomers was extracted for relative quantification and comparison between experimental groups. For co-eluting isobaric peptides (e.g., H3K36me3 and H3K27me2K36me1), the Total Area MS1 was resolved using their unique MS2 fragment ions. The average ratio of analogous ions (e.g., y7 vs. y7) was used to calculate the respective contribution of the precursors to the isobaric MS1 peak.

Relative abundances (percentages) were calculated as follows, using H3K18 acetylation as an example (where “ac” denotes acetylation and “un” denotes unmodified):

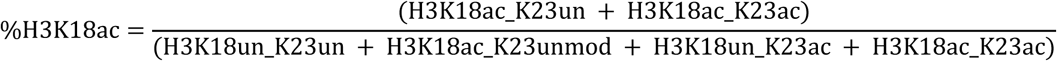

### Quantitative ChIP-seq

U2OS/shUb/HA-Ub(WT) and U2OS/shUb/HA-Ub(K29R) cells were seeded in a 15 cm plate (3 plates per cell line). Cells were treated or not with DOX for 72 h. Fixation buffer A (truChIP Chromatin Shearing Kit, Covaris) was added to each plate, followed by fresh formaldehyde to a final concentration of 1% for fixation at room temperature with constant movement for 10 min. Glycine was then added to quench the reaction for 5 min at room temperature with constant movement. The fixed cells were washed with ice-cold PBS, collected and centrifuged for 5 min at 500g at 4 °C, transferred to Eppendorf tubes, and centrifuged again for an additional 5 min at 500g at 4 °C. The cell pellets were snap-frozen in liquid nitrogen and stored at -80 °C until lysis.

Nuclei isolation was performed on fixed cells using the truChIP Chromatin Shearing Kit (Covaris) according to the manufacturer’s instructions. Cells were sonicated using a Covaris E220 Evolution with settings: 5% duty cycle, 200 cycles/burst, 20 min processing time, 7 °C bath temperature, and full water level. Sonicated chromatin was centrifuged at 14,000 rpm at 4 °C for 10 min, and the supernatant was retained. Drosophila S2 cells were processed in parallel using the same procedure. For each ChIP reaction, 30 μg of U2OS chromatin was mixed with 0.5% sonicated Drosophila S2 chromatin and diluted up to 500 μl with dialysis buffer (4% glycerol; 10 mM Tris-HCl, pH 8.0; 1 mM EDTA; 0.5 mM EGTA) and 400 μl of incubation buffer (2.5% Triton X-100; 0.25% sodium deoxycholate; 0.25% SDS; 0.35 M NaCl; 10 mM Tris-HCl, pH 8.0) supplemented with leupeptin, aprotinin, pepstatin, and PMSF. Chromatin was pre-cleared with Protein A/G Agarose beads (Thermo) for 1 h at 4 °C. After pre-clearing, 10 μl were set aside as input, and the remaining chromatin was incubated overnight at 4 °C with 2 μl H3K9me3 antibody. Protein A/G Agarose beads were pre-blocked with 1 mg/ml BSA in RIPA buffer overnight at 4 °C and then incubated for 3 h with the chromatin-antibody mix.

ChIPs were washed three times in ice-cold RIPA buffer (140 mM NaCl; 10 mM Tris-HCl, pH 8.0; 1 mM EDTA; 1% Triton X-100; 0.1% SDS; 0.1% sodium deoxycholate; 1 mM PMSF), three times in RIPA buffer with 0.5 M NaCl, once in LiCl buffer (250 mM LiCl; 10 mM Tris-HCl, pH 8.0; 1 mM EDTA; 0.5% igepal CA-630; 0.5% sodium deoxycholate), and twice in TE (10 mM Tris-HCl, pH 8.0; 1 mM EDTA). Inputs and washed beads were incubated with 50 μg RNase A (Sigma) for 30 min at 37 °C, followed by the addition of SDS and NaCl to final concentrations of 0.5% and 100 mM, respectively. Samples were incubated with proteinase K (10 μg) for 10 h at 37 °C, followed by 6 h at 65 °C for de-crosslinking. DNA was purified using the MinElute PCR purification kit (QIAGEN) and quantified using a Qubit fluorometer.

The input material and immunoprecipitated DNA were prepared for sequencing using the KAPA Hyperprep kit protocol (Roche) with Illumina-compatible indexed adapters (IDT). The DNA underwent end repair, A-tailing, and amplification, with 6-7 PCR cycles. DNA fragments between 200-700 bp were selected using Agencourt AMPure XP beads (Beckman Coulter) and sequenced paired-end on a NextSeq 2000 (Illumina).

Reads underwent filtering for adapters and low quality using cutadapt (v.4.4). Subsequently, reads were aligned to a hybrid mouse (mm10) and fly (dm6) genome using bowtie2 (v.2.3.4). Duplicate reads were then removed with umi-tools (v.1.1.4), and ENCODE hg38 blacklist regions were masked. Only reads with a mapping quality above 30 were retained for subsequent downstream analyses. QC stats were produced from the bam files according to the ENCODE ChIP-seq pipeline (v.2.2.1).

For spike-in normalization, downsampling factors were calculated for each sample as per ^63^. These factors were determined based on the ratio of ChIP dm6 reads to Input dm6 reads, normalized by the corresponding Input mm10 reads:

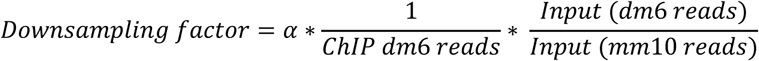

The largest downsampling factor for each protein was set to 1, and other factors were adjusted accordingly. To obtain reference-adjusted reads per-million (RRPMs), the total number of unique reads (uniquely and multi-mapping) were divided by their corresponding downsampling factors.

Visualization tracks for ChIP-seq data were generated using Seqmonk (v.1.47.1), with 5kb non-overlapping bins quantified as indicated for each figure.

Custom R scripts were utilized for creating Hilbert curves, and box plots. These scripts were based on various packages including HilbertCurve and ggplot2.

### Survival analysis

Kaplan-Meier survival analyses were performed using two online tools. Gene Expression Profiling Interactive Analysis (GEPIA2) was used to assess disease free survival stratified by SUV39H1 and TRIP12 expression across 33 cancer types ^36^. Log-rank test was used for hypothesis evaluation. Group cut-off was set at median and a confidence interval of 95% was used. Hazard ratios were calculated following the Cox proportional-hazard model.

Kaplan-Meier Plotter was used to assess overall survival and first progression survival in NSCLC stratified on expression of SUV39H1 and TRIP12 ^38^. Patient cut-off wat set at median. Univariate analysis performing Cox regression to compute hazard ratios and p-value.

### Academia Sinica LUAD-100 Proteome Study

Data from the LUAD-100 cohort were downloaded from the Proteomics Data Commons and processed using the following files: Academia_Sinica_LUAD100_Proteome_v2.tmt10.tsv for proteomics, Academia_Sinica_LUAD100_Proteome_v2.summary.tsv for spectral counts, and Academia_Sinica_LUAD100_Proteome_v2.peptides.tsv for peptide-level information.

Clinical data were retrieved from Academia Sinica LUAD-100_Clinical.xlsx . To advance the proteogenomic understanding of LUAD, the CPTAC program analyzed 111 tumors, of which 102 were paired with adjacent normal tissue samples, subjecting them to global proteome and phosphoproteome analysis. An optimized mass spectrometry workflow using tandem mass tags (TMT-10) was employed ^64^. The LUAD discovery cohort’s proteome, phosphoproteome, and acetylome data are available along with peptide spectrum matches (PSMs) and protein summary reports from the common data analysis pipeline (CDAP) ^65^.

RNA sequencing data for the lung adenocarcinoma (LUAD) cohort was obtained from the Morpheus platform hosted by the Broad Institute. The specific files utilized were for the breast cancer RNA-seq data: rna_seq_breast.gct and for the LUAD RNA-seq data: rna_seq_luad.gct.

Data analysis was conducted using Python, with the primary libraries being pandas for data manipulation, seaborn for visualization, and matplotlib for plotting. Proteomics data were preprocessed by filtering for protein-coding genes and normalizing log-transformed reporter ion intensity values (Log2Ratio). Data from the Academia_Sinica_LUAD100_Proteome_v2.tmt10.tsv file were used, focusing on the expression data across 94 samples. Transcriptomic data were processed by extracting expression values (tpm, fpkm) averaged across the respective aliquots for LUAD cohorts. Expression data was further filtered to focus on protein-coding genes, and subsequent analyses were performed on normalized expression values.

Violin plots were created to visualize the distribution of expression levels for the TRIP12 and SUV39H1 genes across different conditions. These conditions were stratified based on median expression levels of TRIP12, separating the data into “low TRIP12” and “high TRIP12” groups for both proteomics and transcriptomics datasets. For each gene and condition, violin plots were generated using seaborn to compare expression levels across the two groups. This was done separately for proteomics and transcriptomics datasets for both the breast cancer and LUAD cohorts. Additionally, heatmaps were generated using the correlation matrices of the expression data to explore potential relationships between genes. All scripts and code used in this analysis are provided as a Jupyter notebook, which can be accessed along with the raw data files. The notebook includes step-by-step instructions for reproducing the analysis, along with detailed comments to facilitate understanding.

### Sustainability statement

All experimental procedures have been conducted according to the highest sustainability practices as part of the Laboratory Efficiency Assessment Framework (LEAF).

## Acknowledgements

We thank Zhijian Chen, Matthias Altmeyer, Minglei Zhao, Daniel Durocher and Jakob Nilsson for providing reagents, members of the Mailand lab for helpful discussions, and Martin Möckel and the IMB Mainz protein production facility for purification of OtUBD and LotA-N proteins. This work was supported by grants from Novo Nordisk Foundation (grant no. NNF14CC0001), Lundbeck Foundation (grant no. R223-2016-281), Independent Research Fund Denmark (grant no. 9040-00038B), European Union’s Horizon 2020 research and innovation program (Marie-Skłodowska-Curie grant agreement no. 860517 (UBIMOTIF) and no. 846795) and Danish Cancer Society (grant no. R316-A19079).

## Author Contributions Statement

Conceptualization: J.A.G., R.F.S. and N.M.; Methodology: J.A.G., K.W., F.C., D.T., M.V.-A., V.S., and R.F.S.; Investigation: J.A.G., J.W., C.G., E.I., N.R.-G., M.K., A.S.R., K.M., A.I., S.S., A.M., and R.F.S.; Writing – Original Draft: J.A.G., R.F.S. and N.M.; Writing – Review and Editing: All authors; Supervision: L.J.J., A.G., P.B., R.F.S. and N.M.; Project Administration: N.M.; Funding Acquisition: N.M.

---

**Figure S1.**

**Extended data related to Figure 1.**

**A.** Immunoblot (IB) analysis of U2OS/shUb and derivative U2OS/shUb/HA-Ub cell lines treated or not with DOX for the indicated times.

**B.** U2OS/shUb and derivative replacement cell lines were treated or not with DOX and mRNA levels were analyzed by RT-qPCR. Primers to GAPDH were used as a normalization control (data are technical duplicates of two independent experiments).

**C.** Immunoblot analysis of Ub replacement cell lines treated or not with the BRD4 PROTAC MZ1 (500 nM) overnight.

**D.** Abundance of K6 diGly peptide after OtUBD (total Ub) enrichment of indicated DOX-treated Ub replacement cell lines treated exposed to UV-A and S4U as indicated.

**E.** QIBC analysis (left) and immunoblot analysis (right) of DOX-treated Ub replacement cell lines using the indicated linkage-specific antibodies or binders (thick dashed-lines, median; dotted-lines, quartiles). Data is a single representative replicate from 3 independent experiments.

**F.** Cell cycle analysis of indicated Ub replacement cell lines by QIBC. All Ub replacement cell lines were treated with DOX for 72 h, except Ub(K48R) cells that were treated for 48 h. Upper panel shows cell cycle fraction (mean±s.e.m.; *n*=4 independent experiments; >1000 cells analyzed per condition). Lower panel shows mitotic fraction as determined by QIBC of H3-pS10 immunostaining (mean±s.e.m.; *n*=7 independent experiments; >1000 cells analyzed per condition).

**Figure S2.**

**Extended data related to Figure 2.**

**A.** Heatmap comparison of whole proteome analyses of individual replicates from indicated Ub replacement cell lines treated with DOX for 72 h (48 or 72 h in the case of Ub(K48R cells). See **Table S1** for full results.

**B.** Principal component analysis clustering of samples analyzed by MS after enrichment of HA-Ub conjugates from Ub replacement cell lines under denaturing conditions (**Figure 2A-H**).

**Figure S3.**

**Extended data related to Figure 2**

GO term analysis of cellular compartments enriched among proteins showing significantly up- or downregulated ubiquitylation in Ub(K-to-R)-replaced cells relative to Ub(WT)-replaced cells (**Figure 2B-H**).

**Figure S4.**

**Extended data related to Figure 3**

**A**,**B**. Immunoblot analysis of U2OS/shUb, U2OS/shUb/HA-Ub(WT) and U2OS/shUb/HA-Ub(K29R) cell lines treated or not with DOX for the indicated times. Where indicated, cells were treated with Ub E1 inhibitor (E1i) for 1 h.

**C.** QIBC analysis of DOX-treated Ub(WT) and Ub(K29R) replacement cell lines treated or not with the transcription inhibitor Actinomycin D (ActD) or the protein synthesis inhibitor cycloheximide (CHX) where indicated and stained for nascent RNA (upper) or nascent protein (lower panel) by detection of incorporated EU and AHA, respectively (thick lines, median; dotted-lines, quartiles). Data is a single representative replicate from 3 independent experiments.

**D.** U2OS cells or a derivative cell line stably expressing GFP-RNF168 were treated or not with Ub E1i, subjected to GFP pulldown under denaturing conditions and immunoblotted with indicated antibodies.

**E.** DOX-treated Ub(WT) and Ub(K29R) replacement cell lines transfected with FLAG-SUV39H1 expression construct were subjected to FLAG IP under denaturing conditions and immunoblotted with indicated antibodies.

**Figure S5.**

**Extended data related to Figure 4**

**A.** QIBC analysis of Ub(WT) and Ub(K29R) replacement cell lines treated with DOX for the indicated times and immunostained with sAB-K29 (thick dashed line, median; dotted lines, quartiles). Data is a single representative replicate from 3 independent experiments (>1000 cells analyzed per sample).

**B.** QIBC analysis of HeLa cells transfected with indicated siRNAs (thick line, median; dotted lines, quartiles). Data is a single representative replicate from 3 independent experiments (>1000 cells analyzed per sample).

**C.** Immunoblot of HeLa cells transfected with indicated siRNAs.

**D.** Immunoblot analysis of U2OS cells sequentially transfected with indicated siRNA and siRNA-resistant GFP-TRIP12 expression plasmids.

**E.** Representative images of individual cell from data shown in **Figure 4D**. Scale bar, 25 µm.

**F.** DOX-treated Ub(WT) and Ub(K29R) replacement cell lines were subjected to HA IP under denaturing conditions and immunoblotted with indicated antibodies.

**G.** Immunoblot analysis of SUV39H1 in U2OS cells treated or not with cycloheximide (CHX) for the indicated times in the absence or presence of the PARP1 inhibitor Olaparib.

**H.** U2OS/GFP-SUV39H1 cells transfected with siRNAs and treated with Ub E1 inhibitor where indicated were subjected to GFP pulldown under denaturing conditions and immunoblotted with indicated antibodies.

**I.** DOX-treated Ub(WT) and Ub(K29R) replacement cell lines were subjected to SUV39H1 IP and immunoblotted with indicated antibodies.

**J.** Immunoblot analysis of DOX-treated Ub(WT) and Ub(K29R) replacement cell lines treated or not with NEDDylation inhibitor for 4 h.

**K.** As in (H), except that cells were treated or not with NEDDi for 4 h.

**L.** QIBC analysis of U2OS/GFP-SUV39H1 cells transfected or not with HA-TRABID expression construct. Data shown is fold change of mean of average nuclear GFP intensity relative to untransfected cells (*n*=3 independent experiments; *p<0.05, two-tailed t-test).

**M.** Immunoblot analysis of indicated cell lines transfected with non-targeting control (CTRL) or TRABID siRNAs.

**N.** Representative images of U2OS cells transfected with indicated siRNAs and immunostained with sAB-K29.

**Figure S6.**

**Extended data related to Figure 5**

**A.** Immunoblot analysis of soluble and chromatin-enriched fractions of Ub(WT) and Ub(K29R) replacement cell lines treated or not with DOX.

**B.** Hilbert analysis depicting the spatial localization of H3K9me3 across chromosome 11 in DOX-treated Ub(WT) and Ub(K29R) replacement cell lines. Overlap panel depicts signal from Ub(K29R) cells over genes on chromosome 11.

