## Supplementary figures and images for "Targeted disruption of linkage-specific ubiquitylation reveals a key role of K29-linked ubiquitylation in epigenome integrity"

### Supplemental Figure S1-S6

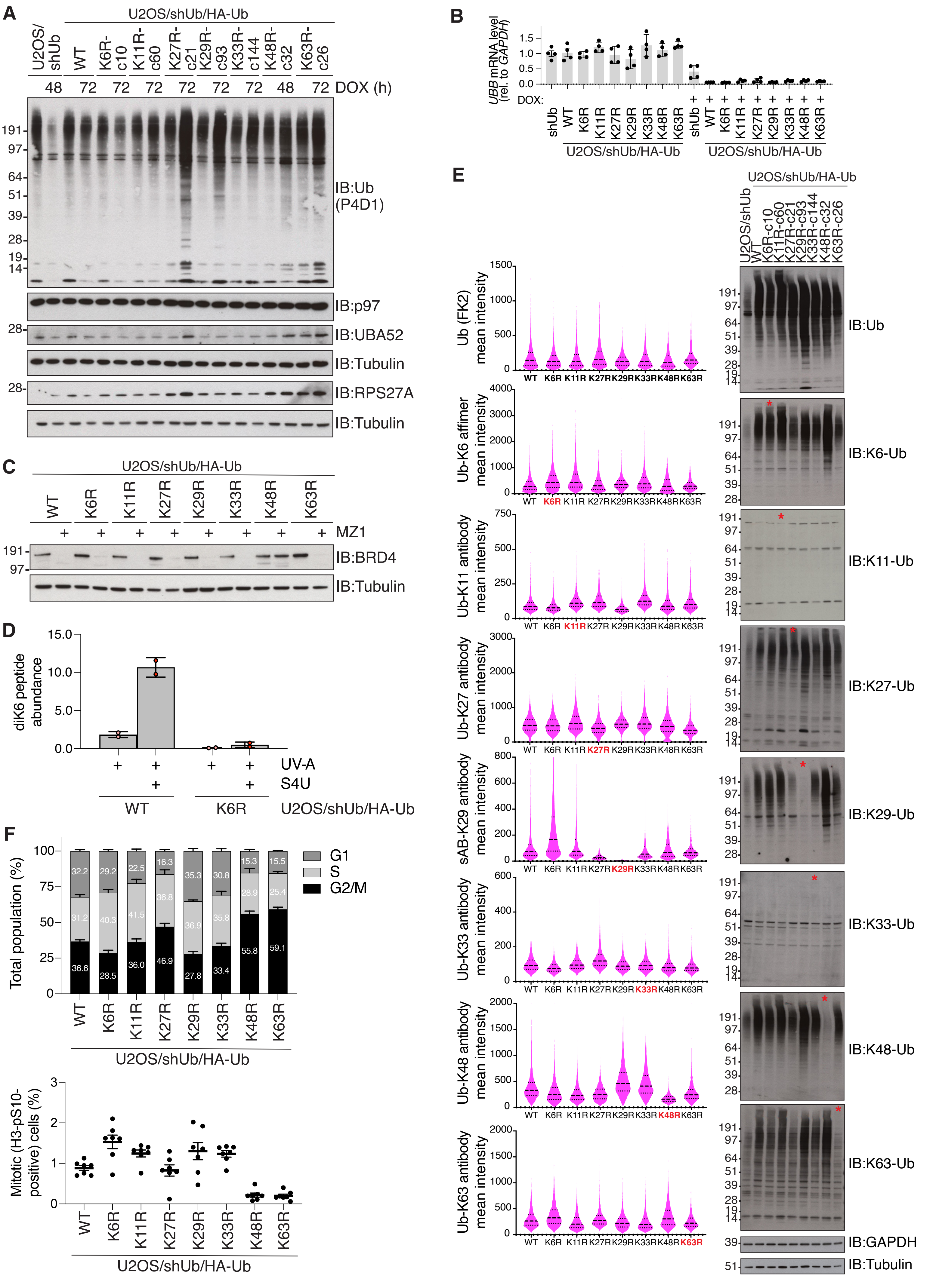

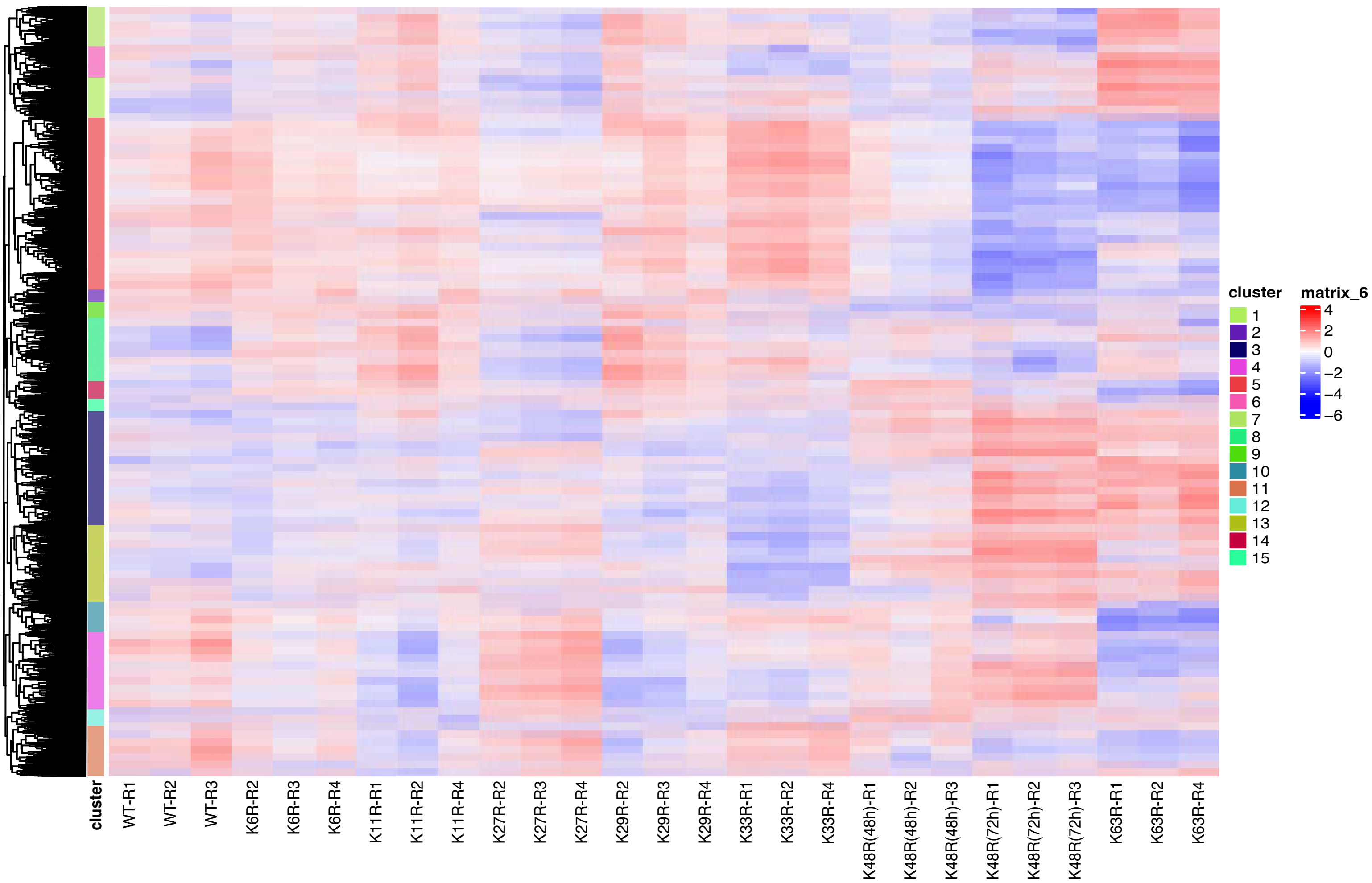

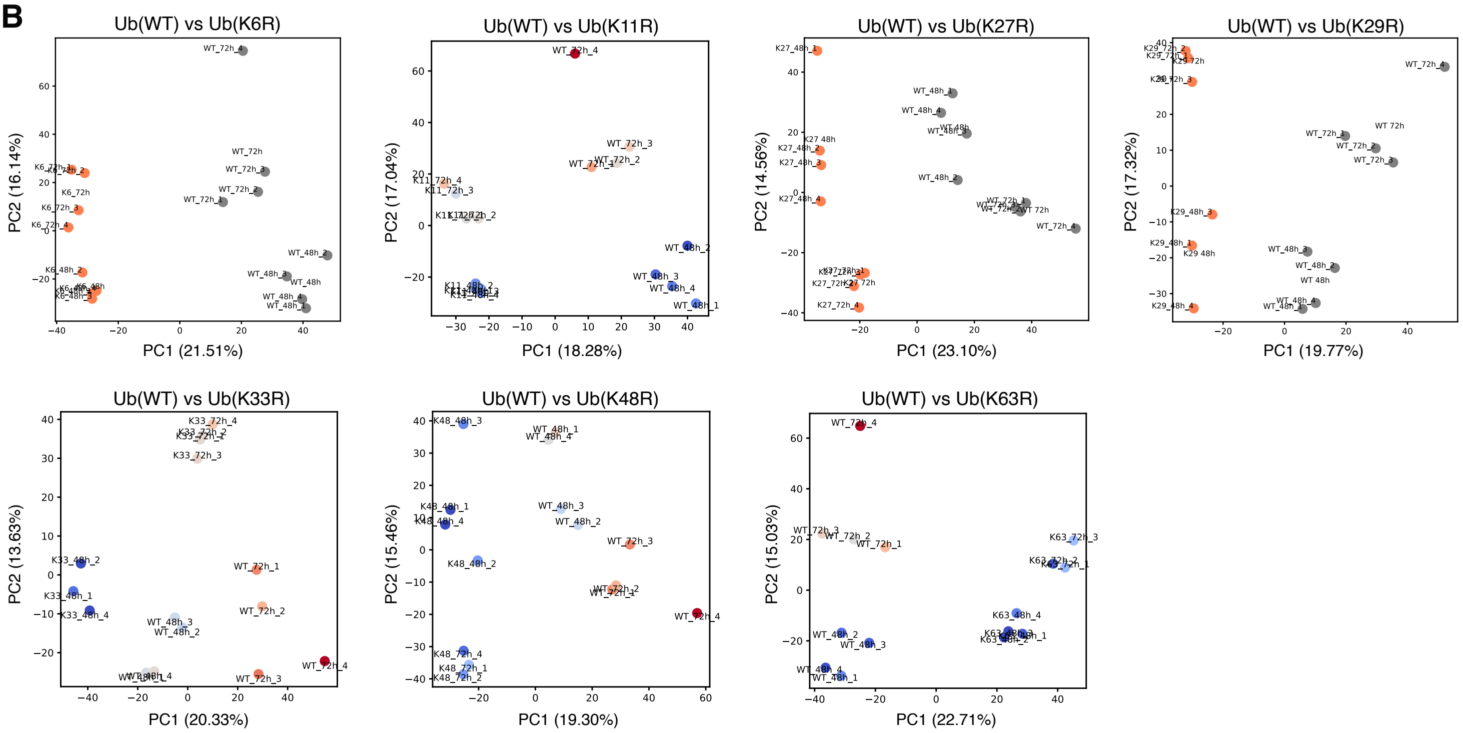

Arroyo-Gomez et al. Fig. S3

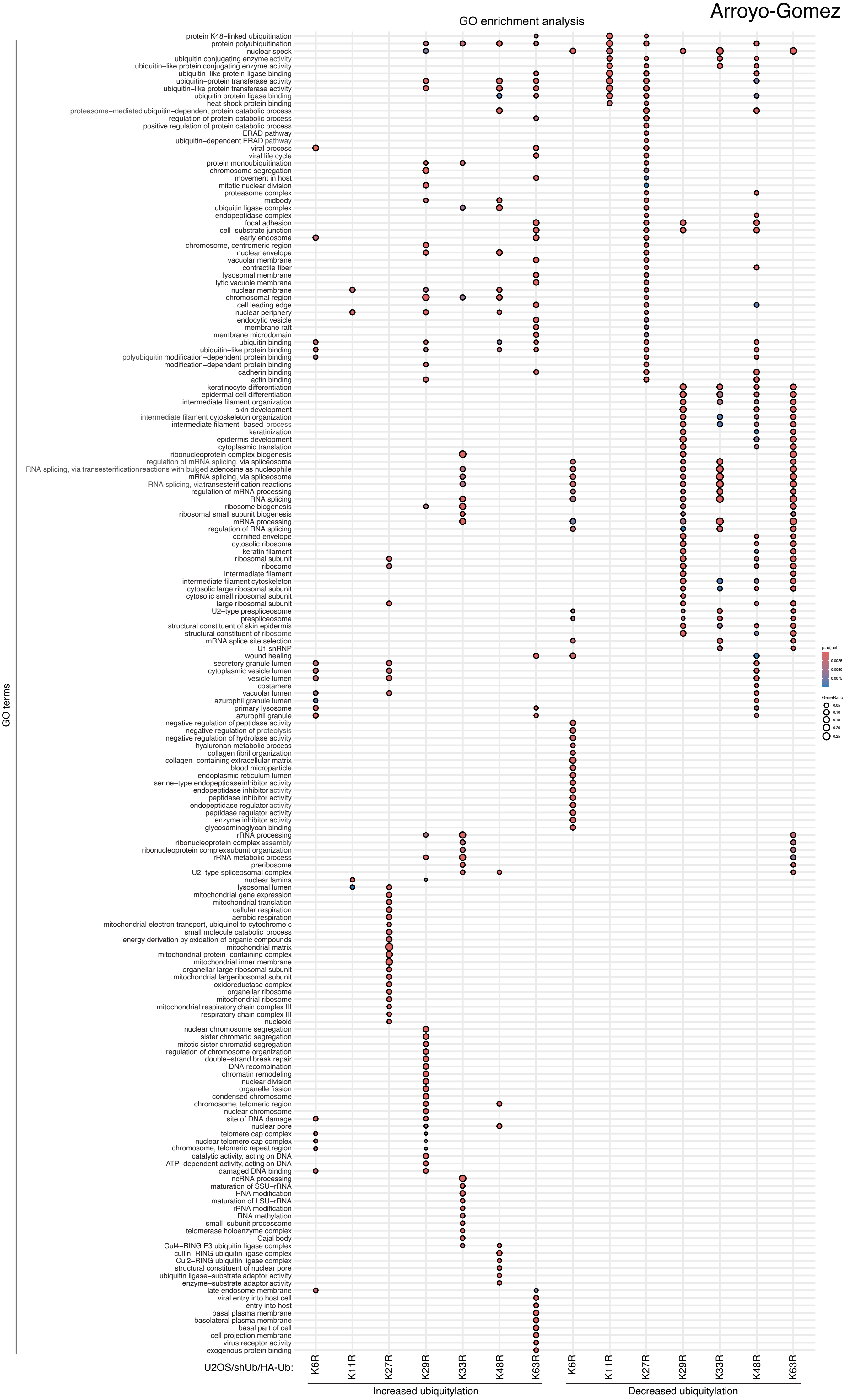

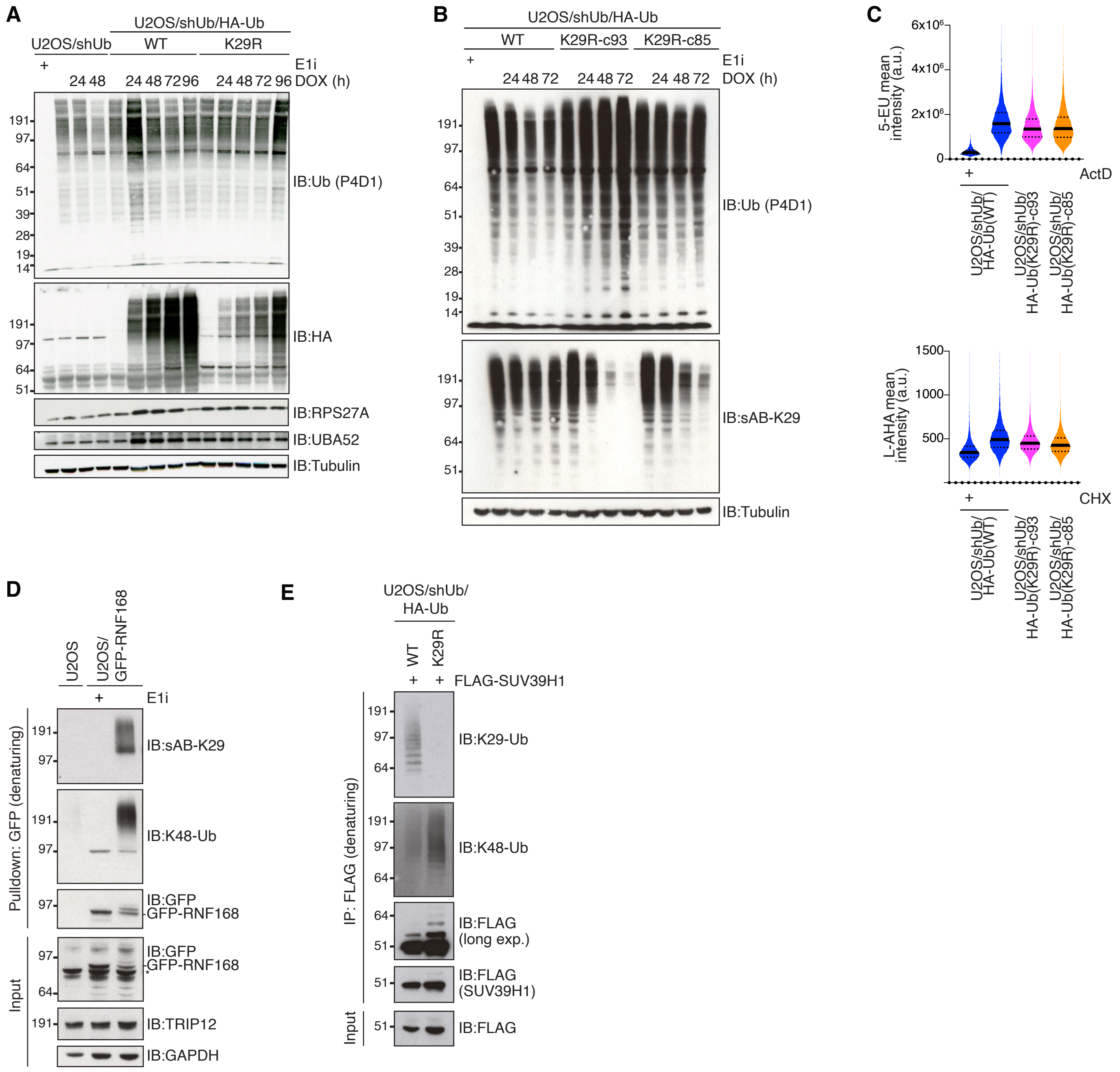

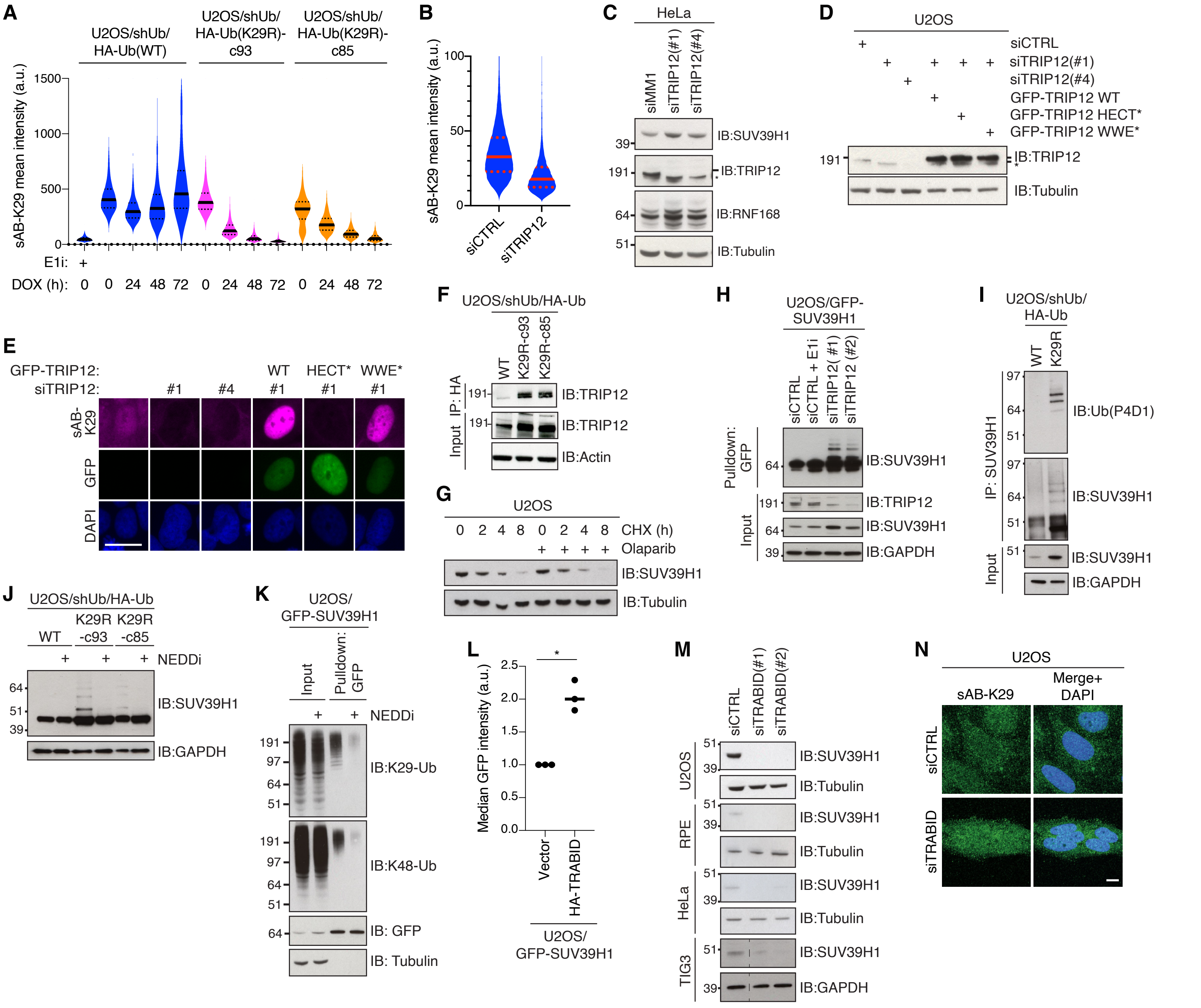

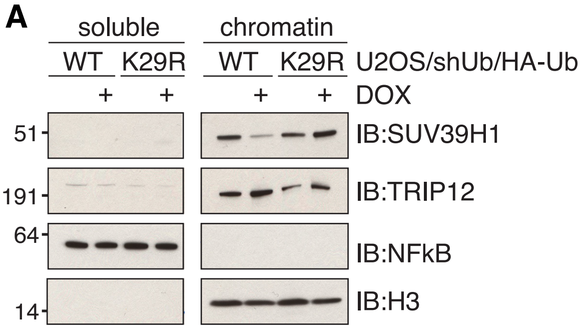

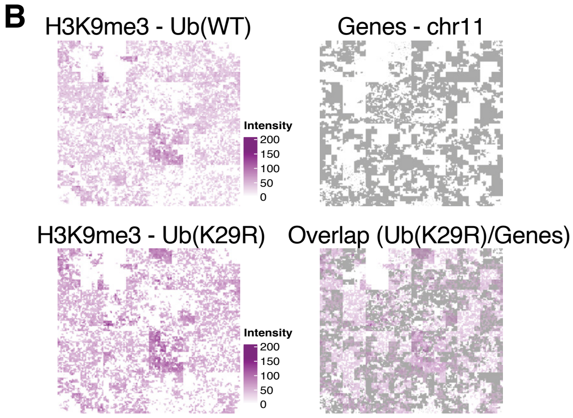
